# Genetic variation at transcription factor binding sites largely explains phenotypic heritability in maize

**DOI:** 10.1101/2023.08.08.551183

**Authors:** Julia Engelhorn, Samantha J. Snodgrass, Amelie Kok, Arun S. Seetharam, Michael Schneider, Tatjana Kiwit, Ayush Singh, Michael Banf, Merritt Khaipho-Burch, Daniel E. Runcie, Victor A. Sanchez-Camargo, J. Vladimir Torres-Rodriguez, Guangchao Sun, Maike Stam, Fabio Fiorani, Sebastian Beier, James C. Schnable, Hank W. Bass, Matthew B. Hufford, Benjamin Stich, Wolf B. Frommer, Jeffrey Ross-Ibarra, Thomas Hartwig

## Abstract

Comprehensive maps of functional variation at transcription factor (TF) binding sites (*cis*-elements) are crucial for elucidating how genotype shapes phenotype. Here we report the construction of a pan-cistrome of the maize leaf under well-watered and drought conditions. We quantified haplotype-specific TF footprints across a pan-genome of 25 maize hybrids and mapped over two-hundred thousand genetic variants (termed binding-QTL) linked to *cis*-element occupancy. Three lines of evidence support the functional significance of binding-QTL: i) they coincide with numerous known causative loci that regulate traits, including *VGT1*, *Trehalase1*, and the MITE transposon near *ZmNAC111* under drought; ii) their footprint bias is mirrored between inbred parents and by ChIP-seq; iii) partitioning genetic variation across genomic regions demonstrates that binding-QTL capture the majority of heritable trait variation across ∼70% of 143 phenotypes. Our study provides a promising approach to make previously hidden *cis*-variation more accessible for genetic studies and multi-target engineering of complex traits.

## Background

Over the past two decades, genome-wide association studies (GWAS) transformed our understanding of the inheritance of many complex traits in important agricultural crops such as maize. Several studies estimated that non-coding variation accounts for about 50% of the additive genetic variance underlying phenotypic diversity in plants (Chia *et al*. 2012; Rodgers-Melnick *et al*. 2016; Lorant *et al*. 2020; Song *et al*. 2021). Although identification of functional non-coding variants is advancing with the development of new genomics technologies (Marand *et al*. 2023) it remains challenging to discern functional variants that impact *cis*-elements efficiently and at cistrome, defined as the genome-wide set of *cis*-acting regulatory loci, scale. Knowing which loci to target has become one of the biggest obstacles for trait improvement via targeted genome editing (Wolter and Puchta 2018; Sharon *et al*. 2018; Marand *et al*. 2023). Scalable methods to construct comprehensive cis-element maps are critical for elucidating complex transcriptional networks that underlie growth, behavior, and disease. The potential of *cis*-element maps has been demonstrated by the ENCODE projects which exist for many eukaryotes, including humans. However, genome-wide, high-resolution maps of functional variants are currently lacking in plants (Lane *et al*. 2014). Despite many successes, GWAS generally suffer from insufficient resolution which limits identification of individual causal single nucleotide polymorphisms (SNPs), insertions or deletions (INDELs), and cannot provide independent molecular information on the potential function of variants, requiring laborious follow-up analyses of numerous individual loci (Sharon *et al*. 2018).

An alternative approach to identify functional polymorphisms would be to annotate non-coding variants within a GWAS region based on their association with transcription factor (TF) binding. This approach has considerable potential, as TF activity plays an important role in the regulation of genes and thereby traits, and the affinity of TF binding is mostly determined by specific local sequences (*cis*-elements) (White *et al*. 2013; Levo and Segal 2014). Identifying *cis*-elements for individual TFs via approaches such as ChIP-seq is time-consuming, not strictly quantitative, limited in scope, and often provides relatively low resolution of functional regions. In contrast, MNase-defined cistrome Occupancy Analysis (MOA-seq) identifies putative TF binding sites globally, in a single experiment with relatively high resolution, yielding footprint regions typically <100bp in size (Savadel *et al*. 2021). In maize, MOA-seq identified ∼100,000 occupied loci, including about 70% of the sequences (bp overlap) identified in more than 100 ChIP-seq experiments (Tu *et al*. 2020; Savadel *et al*. 2021). Notably, many of the MOA-footprint regions were previously uncharacterized, with only 35% identified in previous ATAC-seq data while MOA-seq identified 76% of previous ATAC-seq peaks (Savadel *et al*. 2021). Similarly, an analysis of small MNase-defined fragments from *Arabidopsis* seedlings revealed more than 15,000 accessible chromatin regions missed by ATAC- or DNase-seq (Zhao *et al*. 2020).

Here, we quantified haplotype-specific TF footprints across the maize pan-genome with MOA-seq, utilizing F1 hybrids that share a common reference to minimize biological, technical, and *trans*-effect variation between the haplotypes. We defined a maize leaf pan-cistrome and identified ∼210,000 genetic variants linked to haplotype-specific variation in MOA coverage at *cis*-element loci, we term binding quantitative trait loci (bQTL). bQTL explained the majority of heritable trait variation in >70% of the tested traits in the nested association mapping (NAM) panel. Haplotype-specific TF footprints coincided with causative loci known to affect leaf angle, branching, and flowering time traits, and identified more than 3500 drought-response *cis*-regulatory loci, including *ZmTINY* as candidate loci for future smart breeding.

## Results

### Quantification of functional *cis*-variation

To focus on genetic differences affecting TF binding in *cis*, we quantified TF footprints specific to each haplotype in F1 hybrids with a shared reference parent (B73) (Fig. 1a). We applied MOA-seq to nuclei of both B73 (Manosalva Pérez *et al*. 2024), Mo17 – founders of key maize breeding populations whose hybrid has been extensively studied (Springer and Stupar 2007; Pressoir *et al*. 2009; Hartwig *et al*. 2023) – and their F1 hybrid. MOA footprints were determined by mapping sequencing reads to a concatenated hybrid genome, and retaining reads that mapped uniquely or equally once to both haplotypes (Supplemental Table S1, see methods). We detected 327,029 MOA footprints (at a false discovery rate of 5%) with strong correlation across biological replicates (Pearson’s correlation coefficient > 0.95, Additional file 1, Fig. S1). A total of 53,220 genes in the hybrid, representing 67.9% of the F1 coding sequences (5 kb up- and 1 kb downstream of B73 and Mo17 annotated genes; Supplemental Table S2) were flanked by at least one MOA footprint. Furthermore, the footprint peaks harbored 325,933 sequence polymorphisms, we termed MOA polymorphisms (MPs). We analyzed the allelic ratio of MOA reads at SNPs in the F1, designating the B73 allele as the reference and the other allele as the variant. A total of 48,935 allele-specific MOA polymorphisms (AMPs) deviated significantly from the expected 1:1 allelic ratio (binomial test with 1% FDR, allelic ratio controlled by genomic WGS read analysis, see Methods for details, Additional file 1, Fig. S2).

**Fig 1.**
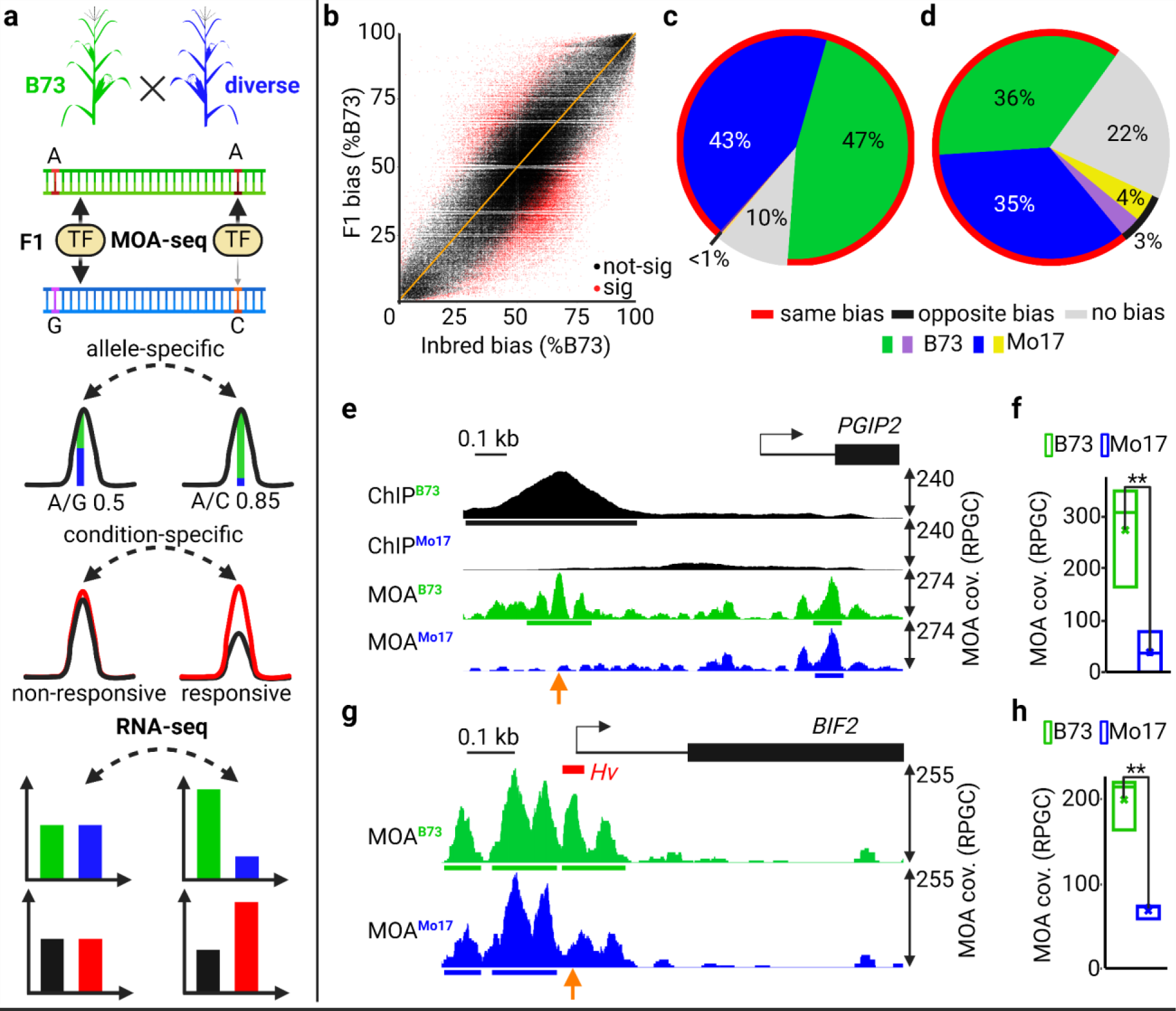
Quantitative *cis*-element occupancy analysis in F1 hybrids. **a**) Haplotype-specific MOA-seq flowchart: 1) Nuclei purified from diverse nested (B73 common mother) F1s are analyzed by MOA-seq, producing small, non-nucleosomal, protein-DNA interaction footprints. 2) SNPs between alleles at MOA sites allow the identification, quantification, and, in a population, association of variants coupled to occupancy of putative *cis*-elements. Allele-specific MOA footprints can be compared between treatments, e.g., well-watered versus drought. 3) Allele-specific mRNA-seq allows further characterization of functional variants associated with gene regulation. **b**) Correlation of haplotype-specific MOA-seq data at all MP loci in nuclei from B73 vs. Mo17 inbreds (X axis) vs. those from the F1 (Y axis) (Pearson correlation 0.78). MPs with (red, p<0.05, expected *trans*) and without (black, expected *cis*) significant differences between F1 and parental alleles. **c**) Genome-wide comparison of allelic bias (50-66.7% to one allele considered no bias, >66.7% considered biased) at B73xMo17 F1 AMP sites to inbred B73 vs. Mo17 data. **d**) Genome-wide directionality analysis, comparing biased SNPs (AMPs) detected by MOA-seq to ChIP-seq data of a single TF BZR1 (Hartwig *et al*. 2023) in the B73xMo17 hybrid. In **c**) and **d**) MOA occupancy was largely consistent (red circle) between either F1 and parents or ChIP-seq, respectively, in showing parental bias towards B73 (green) or Mo17 (blue), with a smaller fraction of MOA sites showing no bias (gray) in ChIP-seq, or bias to the opposite parent/allele (B73 purple and Mo17 yellow). **e-h**) Examples of B73xMo17 MOA-seq (fragment center data, see methods) and allele-specific ChIP-seq. **e**) Average, normalized, allele-specific ChIP-seq of the ZmBZR1 TF (top two rows, black) and MOA-seq (bottom two rows, B73 green, Mo17 blue) shown upstream of *ZmPGIP2*. **g**) Average, normalized, allele-specific differences in MOA coverage upstream of *ZmBIF2* overlap with a known, “hypervariable” (Hv) *cis*-regulatory region. **f, h**) Normalized, average MOA coverage of three biological replicas at orange arrow positions in **e** and **g**. RPGC = number of reads per bin (1 bp) / scaling factor (total number of mapped reads * fragment length) / effective genome size).

The vast majority (88% or 194594/221187) of all MPs showed no significant difference in their allelic bias comparing F1 to B73 vs. Mo17 inbred alleles (black dots Fig. 1b), and nearly 90% (32251/36001) of AMP sites in the B73xMo17 F1 showed bias towards the same allele as in the inbred parents (red line Fig 1c). Fewer than 1% (227/36001) of AMP sites showed bias in the opposite direction. The high concordance between F1 and inbred MOA-seq further establishes the reproducibility of the assay and indicates that the majority of AMPs are coupled to genotypic differences in *cis* at the binding site, rather than resulting from *trans*-acting or *cis*-by-*trans* interaction effects.

To independently validate haplotype-specific, MOA-defined TF footprints in B73xMo17, we compared AMPs to recently published allele-specific ChIP-seq data of the major brassinosteroid TF ZmBZR1 in the same F1 (Hartwig *et al*. 2023). More than 70% of AMPs overlapping with ZmBZR1 binding sites showed allelic bias in the same direction in both studies (red line Fig. 1d). About 20% of AMPs show no bias in the ChIP-seq data, likely due to the lower resolution of haplotype-specific ChIP-seq (∼500 bp fragments compared to ∼65 bp for MOA-seq). Only 8% of AMPs show bias for different alleles than in ChIP-seq, potentially reflecting biological differences in the tissues analyzed (meristem and leaf vs. leaf) or ectopic BZR1 activity due to exogenous brassinosteroid treatment (Hartwig *et al*. 2023).

If haplotype-specific MOA-seq can detect relevant variation in TF binding, we expect it to coincide with potential *cis*-variation in the B73xMo17 F1. We previously reported the allele-specific binding of ZmBZR1 upstream of the TSS of *POLYGALACTURONASE-INHIBITING PROTEIN2* (*ZmPGIP2*, Zm00001eb034870, Fig. 1e) (Hartwig *et al*. 2023). PGIP2 is a cell wall protein and a candidate locus for both northern and southern leaf blight resistance (Balint-Kurti *et al*. 2007). The overlapping MOA and BZR1 ChIP binding peaks were both significantly higher for the B73 allele compared to Mo17, exhibiting a 5-fold (p<0.01) higher MOA coverage (Fig.1e-f). Each of the B73 AMP footprints overlapped with known ZmBZR1 BRRE or G-Box motifs (CGTGTG, CACGTG, and CACGTT, respectively). In contrast, the Mo17 allele contained both a SNP in the BRRE motif as well as a HIP-superfamily helitron insertion between the BRRE and G-Box motifs. In another example, we found a significantly higher (1.96-fold; p=0.016) MOA occupancy for the B73 allele for a region upstream of *BARREN INFLORESCENCE2* (*ZmBIF2,* Zm00001eb031760) (Fig. 1g-h), along with codirectional changes in transcript abundance (Additional file 1, Fig. S3). The MOA-seq AMP footprint overlaps a small 80 bp hypervariable region in the *BIF2* proximal promoter (in Fig. 1g) previously associated to differences between B73 and Mo17 in traits such as tassel branch zone length, plant height, and leaf width and length (Pressoir *et al*. 2009). Together, these examples illustrate the potential of MOA-seq to annotate candidate *cis*-regulatory elements with quantitative chromatin footprint data that connects *cis*-variation to biases in *cis*-element occupancy.

### Defining functional sites in a maize pan-cistrome

To define a leaf pan-cistrome of maize, we analyzed a population of 25 F1 hybrids using haplotype-specific MOA-seq (Fig. 1a,). The hybrid population, created by crossing to 25 inbred lines with high quality genome assemblies (Springer *et al*. 2018; Sun *et al*. 2018; Lin *et al*. 2021; Hufford *et al*. 2021) to the reference genome line B73, represents a diverse set of maize including many of the parents of an important mapping population and several important genetic stocks (Supplemental Table S3). We analyzed allele-specific TF occupancy and mRNA abundance in leaf blades for three biological replicates of each hybrid (Supplemental Tables S1 and S4). By aligning MOA-seq and RNA-seq reads to concatenated dual-reference genomes rather than a single reference, our approach resolves issues of reference bias that confound most allele-specific analyses (Fig. 2a, see methods (Castel *et al*. 2015). We identified an average of 362,000 MOA peaks (at a false discovery rate of 5%) per F1, covering approximately 2% (around 80 Mbp) of each hybrid genome (Additional file 1, Fig. S4). On average 19.9% (14-30%) of MPs showed a significant allelic bias (binomial test, false discovery rate of 1%, Supplemental Table S5) with an overall even split between the parental alleles (51% B73 and 49% diverse parents, Additional file 1, Fig. S5, Supplemental Table S5). It is noteworthy that the average rate of AMPs (19.9%) closely matches allele-specific TF binding sites detected by the gold standard of ChIP-seq for an individual TF (18.3%) (Hartwig *et al*. 2023). In total, AMPs overlapped with 36.1% of all MOA footprint peaks in B73 (Fig. 2b), and plots of the identified MOA peaks and cumulative bp indicate our sample is near saturation and has identified the majority of the B73 cistrome (Additional file 1, Fig. S6).

**Fig 2.**
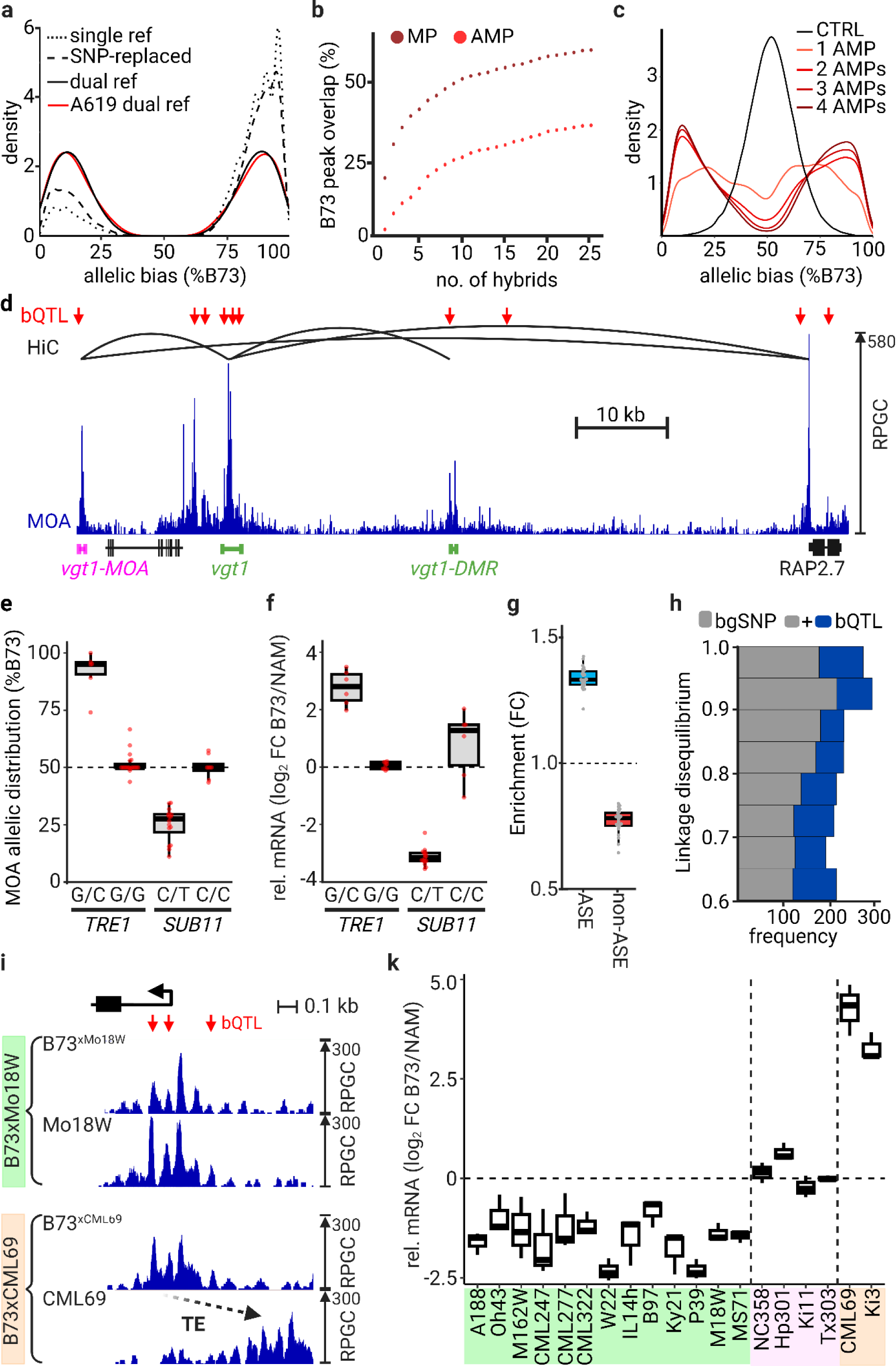
A Pan-cistrome of the maize leaf. **a**) Mapping strategies comparison showing the distribution of allele-specific occupied sites (% binding to B73). B73xHP301 data was analyzed using either only B73 as reference (single ref), a pseudo-genome with B73/HP301 SNPs replaced by Ns (SNP replaced), or our dual-parent mapping strategy using a concatenated B73 x HP301 genome (dual ref). For the A619 F1 (no assembled genome available), our “reference-guided” strategy showed similar AMP-balanced haplotypes without reference bias (A619 dual-ref). **b**) Percent of B73 MOA peak covered by MPs (brown) and AMPs (red) relative to the number of F1s. **c**) Density of mean MOA binding frequencies at SNP positions where at least one, two, three or four lines had AMPs, compared to a control with randomized binding frequencies. **d**) Overview of bQTL (red arrows), MOA coverage (blue), and HiC interaction sites (black lines, HiC from Ricci et *al*., 2019) near the classical flowering repressor *RAP2.7*. MOA bQTL overlap both known enhancers, *vgt1* and *vgt1-DMR* (green), associated with *RAP2.7* expression. An additional bQTL, termed *vgt1-MOA* (magenta), also interacts with *vgt1* and the RAP2-7 promoter. **e**) Allelic distribution (%B73) of MOA reads at bQTL in either the *TRE1* (Zm00001eb021270, bQTL: B73-chr1:80,826,022) or *ZmSUBTILISIN11* (Zm00001eb152020, bQTL B73-chr3:198,733,446) proximal promoters. For *TRE1*, 19 F1s sharing the B73 allele (G/G), were compared to 6 F1s carrying G/C alleles. For *ZmSUBTILISIN11*, 8 F1s sharing the B73 allele (C/C) were compared to 16 F1s with C/T alleles. **f**) Haplotype-specific mRNA counts for *TRE1* and *ZmSUBTILISIN11* grouped by their respective bQTL alleles in (**e**). **g**) ASE and non-ASE genes are significantly more and less enriched for AMPs in their 3 kb promoter upstream of the TSS, respectively. **h**) MOA bQTL are more often in linkage disequilibrium with *cis-*expression QTL, identified in roots of 340 recombinant inbred lines than matched bgSNPs. **i**) Average, normalized MOA coverage for B73 (top) and NAM (bottom) alleles of B73xMo18W and B73xCML69 upstream of phosphoglycerate mutase (*PGM1*, Zm00001eb196320). Compared to B73, the Mo18W allele showed significantly higher MOA occupancy upstream of PGM1 (green panel). In contrast, the CML69 allele showed similar MOA coverage compared to B73, yet peaks were shifted ∼300 bp due to a mite TE insertion (orange panel). **k**) Allele-specific mRNA counts of *PGM1* in the different F1 hybrids. Colors indicate MOA ratios at bQTL: NAM > B73 (green, >60% bias to NAM, non-B73 allele for at least one bQTL), B73 = NAM (purple, %B73 occupancy between 40% to 60%, sharing B73 genotype), and TE insertion haplotype (orange).

We next sought to identify (epi)genetic variants associated with differences in MOA-detected TF occupancy between haplotypes, or binding quantitative trait loci (bQTL), in our population. We first verified that F1s that shared haplotypes at AMP loci also share similar patterns of allelic bias (Fig. 2c), suggesting that our SNPs were in sufficient linkage disequilibrium (LD) with causal differences to perform association analysis. Differences in DNA methylation between parental alleles can affect TF binding affinity (O’Malley *et al*. 2016; Hartwig *et al*. 2023). After validating that DNA methylation differences at AMPs detected between F1 haplotypes were consistent between the parental lines (Additional file 1, Fig. S7), we added previously published methylation data for 24 of our parental lines (Liang *et al*. 2021; Lin *et al*. 2021; Hufford *et al*. 2021). Mixed linear modeling identified 163,395 bQTL significantly (FDR<0.05) associated with variation in MOA-defined *cis*-element occupancy (Supplemental Table S6). SNPs alone explained 45% of the observed differences in MOA coverage, whereas SNPs and DNA methylation combined explained 57% (Additional file 1, Fig. S7). As expected the genome-wide distribution of bQTL was distinct from all SNPs and more closely matched those of TF target sites (Tu *et al*. 2020) and allele-specific ZmBZR1 binding sites (Additional file 1, Fig. S8, (Hartwig *et al*. 2023)).

### bQTL coincide with numerous known, causative regulatory loci

Detailed analyses of regulatory variation for a number of maize genes provide an opportunity to compare bQTL to previously identified causal variation. For example, a bQTL was directly adjacent to the YABBY TF binding site underlying the leaf architecture QTL *Upright Plant Architecture2* (Tian *et al*. 2019, Additional file 1, Fig. S9). bQTL also identified haplotype-specific footprints at a number of flowering time loci, including the causative transposon insertions at *ZmCCT9* and *ZmCCT10* (Huang *et al*. 2018), INDEL-2339 in the promoter of the FT-like *ZmZCN8* (Meng *et al*. 2011), as well as multiple GWAS hits for flowering time (Additional file 1, Fig. S9). In addition to identifying bQTL in both of the known distal regulatory regions*, VGT1* and *VGT1*-DMR, of the key flowering time locus *ZmRAP2.7* (Salvi *et al*. 2007; Castelletti *et al*. 2014; Xu *et al*. 2020), our bQTL analysis identified a novel, third putative enhancer more than 100 kb upstream, which we termed *VGT1-MOA* (Fig. 2d). HiC long-range interaction data (Ricci *et al*. 2019) confirmed that *VGT1-MOA* physically interacts with both *VGT1* and the proximal *ZmRAP2.7* promoter (Fig. 2d).

Transposable element (TE) insertion plays an important role in natural variation of gene regulation (Noshay *et al*. 2021). While the repetitive nature of transposons makes overlapping footprint analysis challenging, we found bQTL at numerous TE-associated *cis*-regulatory regions. In addition to the bQTL that coincide with flowering loci, we observed that bQTL colocalize with the regulatory variation upstream *ZmTB1* and *ZmGT1*, which form a regulatory module involved in bud dormancy and growth repression (Dong *et al*. 2019). bQTL were adjacent to both causal, distal transposon insertions upstream of *ZmTB1* (Studer *et al*. 2011), and coincide with the TE-associated causal regulatory region (prol1.1) upstream of *ZmGT1* (Wills *et al*. 2013) (Additional file 1, Fig. S10).

Our MOA-seq pan-cistrome provides an opportunity to evaluate not only overlap with known trait-associated sites, but how variation at these sites compares to changes in cis-element occupancy. For example, an INDEL in the *TREHALASE1* promoter (*TRE1*) has been associated with both trehalose levels and *TRE1* transcript levels in maize (Wen *et al*. 2018). We observed haplotype-specific footprints, both at the previously reported 8 bp insertion (Wen *et al*. 2018) and an additional SNP 29 bp upstream, which coincided with a bQTL (Additional file 1, Fig. S11).

Notably, while the 8-bp insertion creates a potential ABI motif (TGCCACAC), the *TRE1*-bQTL overlaps with a DOF binding motif (AAAAGGTG). Indeed, previously published ChIP-seq results confirm that the *TRE1*-bQTL site is targeted by ZmDOF17 (Additional file 1, Fig. S11, Tu *et al*. 2020). Furthermore, all alleles (6/6) in our F1 population without the 8 bp insertion and with the non-canonical DOF motif (C instead of G) at the bQTL site, showed concomitant low haplotype-specific MOA signal (strong bias towards B73’s G allele) and *ZmTRE1* mRNA levels (higher B73 mRNA level) (Fig. 2e-f). In another example, *ZmSUBTILISIN11* has been associated with cell wall compositions, peduncle vascular traits, and ABA levels (Cui *et al*. 2023). A previously identified *cis*-expression QTL lead SNP for *ZmSUBTILISIN11* transcript levels (Sun *et al*. 2023) coincided with a bQTL in its proximal promoter and we observed a strong correlation of haplotype-specific MOA footprints at the bQTL and *ZmSUBTILISIN11* transcript levels (Fig. 2e-f).

### Variation in MOA occupancy is correlated to differential transcript accumulation

If variation in MOA coverage accurately captures TF binding affinity, we expect to see associations between haplotype-specific MOA coverage and transcript abundance in our F1s. Indeed, we find that the promoters (within 3 kb upstream of the TSS) of genes with significant haplotype-specific transcripts abundance (p<0.05, see methods) were ∼35% and ∼75% enriched for the presence of AMPs compared to all expressed and non-differentially (p>0.95) expressed genes, respectively, in both well-watered (Fig. 2g) and drought conditions (Supplemental Table S7, Additional file 1, Fig. S12). MOA bQTL were also substantially more likely to be in high linkage disequilibrium (> 0.6) with nearby *cis*-expression QTL in a panel of 340 maize genotypes (Sun *et al*. 2023) than matched background (bg) SNPs (enrichment for intergenic 839/529 (58.6%) bQTL vs. bgSNPs, and 3478/2894 (20.2%) for all bQTL vs. bgSNPs, respectively) (Fig. 2h). These broad patterns are reflected at the level of individual genes as well. For example, all of the NAM parents showing greater MOA occupancy at the two bQTL upstream of phosphoglycerate mutase (*ZmPGM1*, Zm00001eb196320) showed increased abundance of the NAM transcript, whereas F1s with no polymorphism between B73 and NAM showed no difference in the haplotype-specific transcript levels (Fig. 2i-k). Two NAM parents, Ki3 and CML69, showed drastically reduced *ZmPGM1* mRNA levels compared to B73 in both the F1 (Fig. 2k) and inbred lines (Hufford *et al*. 2021), despite similar MOA coverage. The MOA peaks for the Ki3/CML69 haplotypes, however, were shifted by approximately 300 bp compared to B73 due to the insertion of a PIF/Harbinger transposon resulting in hypermethylation of the DNA between the MOA peak and the TSS, a pattern not observed for alleles without the transposon (Fig. 2i-k, Additional file 1, Fig. S13). Taken together, our results show that variation at bQTL correlates with differences in gene regulation.

### Variation in DNA-methylation can predict MOA occupancy

The vast majority of TFs in *Arabidopsis* have been shown, *in vitro*, to have higher binding affinity to hypomethylated DNA (O’Malley *et al*. 2016). We explored this association in our data, focusing on mCG/mCHG variation, as they accounted for >99.8% of methylation differences at MOA sites. Significant DNA methylation differences (following Regulski *et al*. 2013), one allele <10% methylated and the other >70%) overlapped with 14.8% of MPs in the F1s. At AMP loci haplotype-specific mCG/mCHG overlap increased by 2.6-fold (38.1%) and reached more than half (51.5%) for AMPs with a strong haplotype-bias (at least 85% to one allele) (Fig. 3a). We observed a very strong correlation between a higher footprint occupancy and the hypomethylated allele (Fig. 3b) with 98.2% of AMPs showing higher MOA coverage at the hypomethylated allele. Furthermore, nearly half of the remaining 1.8% AMPs biased towards hypermethylation alleles did not display methylation difference immediately surrounding the AMP (10 bp window), suggesting there may be no actual methylation difference at the occupied site despite hypermethylation of the surrounding region (40 bp window) (Additional file 1, Fig. S14). On average, the vast majority of F1s _[MS1]_ that shared differentially methylated alleles (71.2% of these F1s) at a given loci also shared haplotype-specific MOA footprints at that site, significantly more the F1s with shared equally methylated alleles at the same site (42.9% of F1s with equally methylated alleles, Fig. 3c). The observed strong correlations between differential CG and CHG methylation and haplotype-specific MOA occupancy confirm an important role for DNA methylation in determining TF binding in maize.

**Fig 3.**
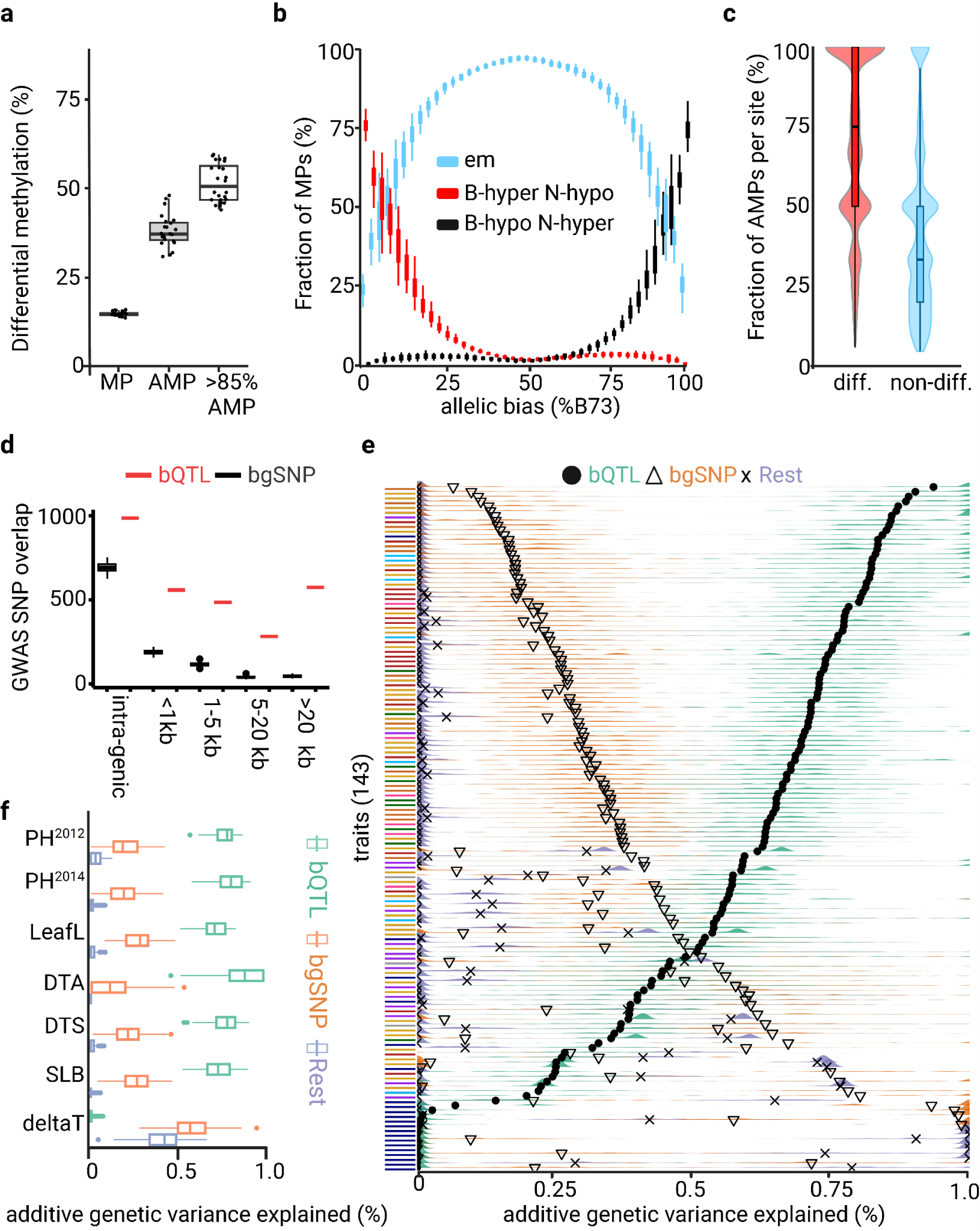
bQTL are linked to variation in DNA methylation and traits. **a**) Genome-wide overlap of differentially methylated (CG and/or CHG) DNA regions with MPs, AMPs, and AMPs that showed a strong (>85%) allelic bias, across the 24 F1s. **b**) Correlation of differentially CG methylated DNA across the allelic bias for MPs in the 24 F1 hybrids. MP methylation categories: equal methylated (em, blue), B73 hyper-vs. NAM hypo- (red), or NAM hypo- vs. B73 hyper- (blue) methylated. **c**) Correlation between MOA footprint bias and differential methylation at loci which varied both in allele-specific footprint occupancy (≥1 F1s with AMP) and CG methylation (≥2 F1s with and without allele-specific methylation difference) between the 24 F1s. Box and violin plots of percentage distribution of F1s with haplotype-specific binding (AMPs) at these positions, partitioned into F1s with either differential allelic CG methylation (red) or equal CG methylation (blue). **d**) Association of ∼42k curated GWAS hits (+/− 100 bp, (Tian *et al*. 2020)) with bQTL or matched bgSNPs at distances ranging from intra-genic to >20 kb to the nearest gene. **e**) Estimated additive genetic variance organized by 143 traits. Colored ridges show the estimated additive genetic variance across 100 permutations for either MOA bQTL (green), bgSNPs (orange), and remaining genome SNPs (purple). Black symbols represent the mean estimated value across permutations. Traits arranged by bQTL mean variance estimates and color coded according to general trait groupings: vitamin E metabolites = navy blue, metabolites = purple, stalk strength = light blue, flowering time = gold, plant architecture = red, disease = green, tassel architecture = pink, ear architecture = orange, misc. = grey. **f**) A subset of traits (y-axis) and their estimated percent additive genetic variance (x-axis) shown as colored boxplots instead of ridges. PH=plant height (Hung *et al*. 2012; Peiffer *et al*. 2014), LeafL=leaf length (Hung *et al*. 2012), DTA=days to anthesis (Hung *et al*. 2012; Peiffer *et al*. 2014), DTS=days to silking (Hung *et al*. 2012), SLB=southern leaf blight (Kump *et al*. 2011; Bian *et al*. 2014), and delta-tocopherol concentration = vitamin E biosynthesis (Diepenbrock *et al*. 2017).

### MOA bQTL explain a large portion of heritable variation

Regulatory variation is thought to underlie a significant proportion of phenotypic variation in maize (Wallace *et al*. 2014). To assess the relationship between MOA bQTL and complex trait variation, we first quantified the enrichment of bQTL surrounding GWAS hits (lead SNP +/- 100 bp) across two curated datasets of 41 and 279 traits (Wallace *et al*. 2014; Tian *et al*. 2020). In both cases, MOA bQTL were approximately twofold enriched for co-localization with GWAS hits compared to matched bgSNPs with similar allele frequency and distance to the nearest gene (100 permutations, see methods, Additional file 1, Fig. S15). This enrichment remained stable as a function of distance to the nearest gene, indicating comparable efficacy of bQTL to mark functionally significant loci genome-wide (Fig. 3d). To explore the degree to which bQTL can more broadly capture the genetic variation underlying phenotypic diversity, we partitioned heritable trait variance for 143 traits in the NAM population (see methods, (Rodgers-Melnick *et al*. 2016; Hartwig *et al*. 2023)). We modeled additive genetic variation for traits as a function of genomic relatedness matrices estimated from bQTL, matched bgSNPs, and SNPs from the rest of the genome. Variances estimated this way for several trait data sets simulated from our observed matrices accurately reflected the proportional contributions of each SNP set (Additional file 1, Fig. S16). Across a large majority of phenotypes in the NAM panel (101 of 143 or ∼71%), bQTL explained the majority of the total additive genetic variance captured by SNPs (Fig. 3e, Additional file 1, Fig. S17, Supplemental Table S8). Consistent with previous findings that open chromatin and TF binding play a key role in trait variation (Rodgers-Melnick *et al*. 2016; Hartwig *et al*. 2023), our matched bgSNPs (matched for similar allele frequency and distance to the nearest gene) often accounted for more additive genetic variation than SNPs from the rest of the genome (123 out of 143 traits), but MOA bQTL also outperformed bgSNPs for most traits (75.7%, 106 traits; Fig. 3e) and randomly sampled SNPs within MOA peaks (Additional file 1, Fig. S18). Traits where bQTL explained the largest portion of genetic variance included plant height, leaf size or shape, and disease-resistance, while almost all traits related to e.g. vitamin E production were best explained by the bQTL-matched bgSNPs or the remaining SNPs from the rest of the genome (Fig. 3f), likely because of the oligogenic nature of the vitamin E traits and that MOA bQTL identified in leaf tissue may not be representative of regulatory patterns in genes specifically expressed in kernels (Diepenbrock *et al*. 2017).

### Characterization of a drought-responsive cistrome

We next compared the morphological and molecular response of our F1 population under well-watered (WW) and drought (DS) conditions to evaluate variation in MOA occupancy induced by a response to changes in environmental conditions. We observed diverse responses in the V4-stage plants subjected to 86h of withholding watering, with a reduction of relative leaf water content ranging from 10-32% and remaining soil water content ranging from 6.3 % to 25.6 %, depending on the F1 (Fig. 4a-b, Additional file 1, Fig. S19 & S20). To evaluate drought-induced differences in *cis*-element regulation, we performed haplotype-specific MOA- and RNA-seq. The number of MOA peaks showing significant (p<0.05) drought-induced increases or decreases in occupancy varied substantially among F1s, ranging from around 23k to 83k and 37k to 134k, respectively (Supplemental Table S9). Local association mapping identified 126,008 DS-bQTL under drought conditions (for a combined total of 208,979 unique bQTL) (Supplemental Table S10). To identify a set of candidate drought-response loci, we selected bQTL with drought-responsive occupancy near genes (5 kb up- / 1 kb downstream) that displayed both haplotype-specific and drought-responsive transcript accumulation, resulting in 995 (417 genes) and 3201 (1194 genes) bQTL with increased and decreased occupancy, respectively (referred to as DS-bQTL). Further integration with GWAS and *cis*-expression QTL hits for drought-response traits (Li *et al*. 2016; Shikha *et al*. 2017; Wu *et al*. 2021) resulted in high-confidence candidates (Supplemental Table S11). Notably, the candidate list included known drought-response variation, such as in the proximal promoter of the “NAM, ATAF, and CUC” (NAC) TF *NAC111*. A haplotype-specific MOA footprint identified by a DS-bQTL overlapped with the causative 84 bp MITE transposon insertion site, which reduces both *ZmNAC111* expression and drought tolerance in maize seedlings, likely through RNA-directed DNA methylation (Mao *et al*. 2015) (Additional file 1, Fig. S21). A DS-bQTL also located within the previously discovered 119 bp proximal promoter fragment required for the drought-response of *SULFITE OXIDASE1* (*ZmSO*, Zm00001eb036560), a gene linked to the ABA-response and drought-tolerance of maize seedlings (Xu *et al*. 2019). Although none of the haplotypes in our population contained the putative Myb-binding site (CAGTTG) previously linked to the drought-response in the 119 bp *ZmSO* promoter (Xu *et al*. 2019), we nonetheless found a strong correlation between increased MOA occupancy for the C allele at the bQTL and elevated *ZmSO*^B73^ transcript levels both under WW and DS conditions (Fig. 4c-d). We also observed a strong correlation between MOA occupancy and drought-induced transcript levels at DS-bQTL in the proximal promoter of the maize homolog of aquaporin *BETA-TONOPLAST INTRINSIC PROTEIN 3* (*ZmTIP3d*, Zm00001eb076690, Fig. 4e-f) which has been linked to drought-response in various plants (Johansson *et al*. 2000).

**Fig 4.**
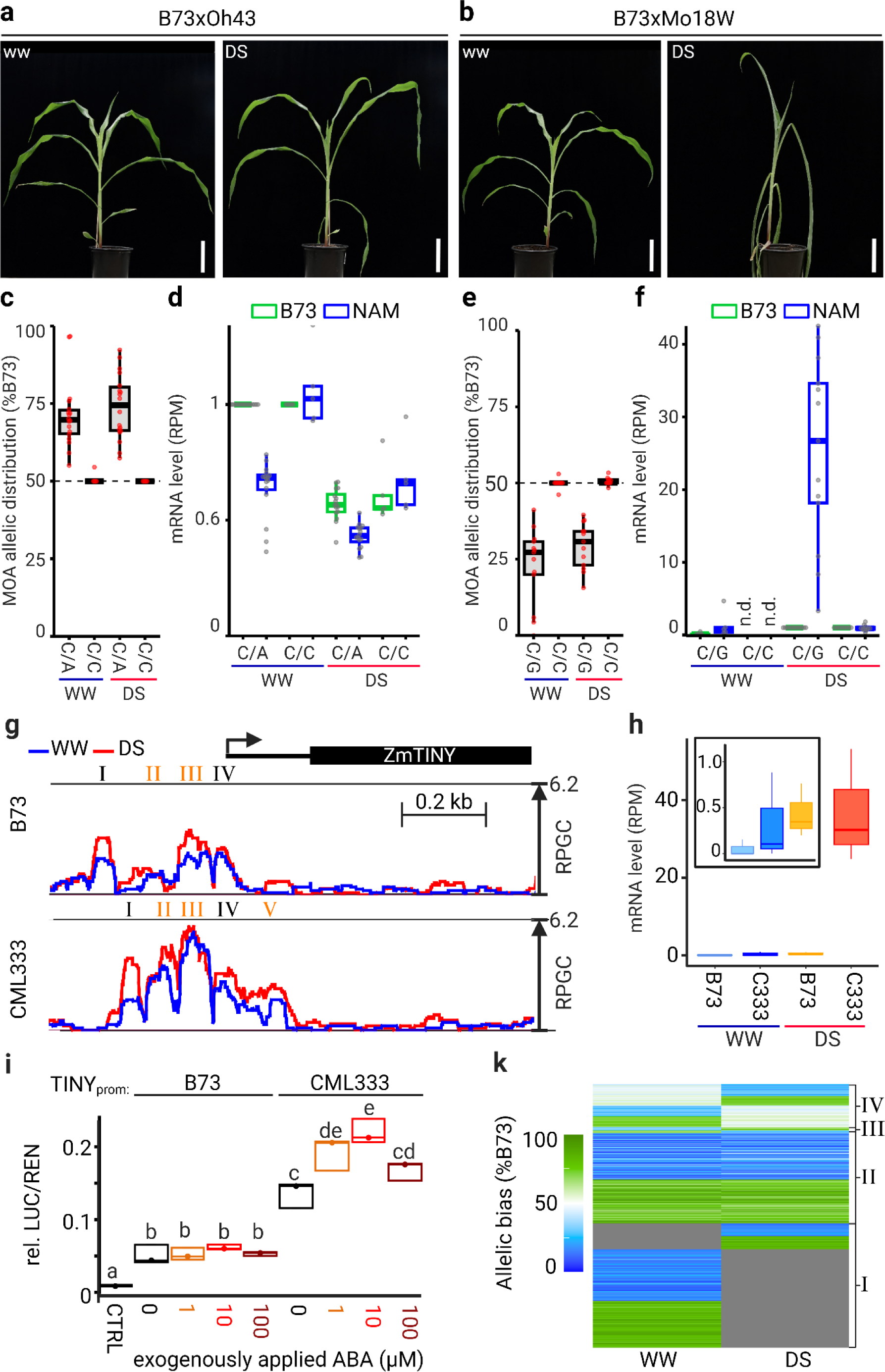
MOA-seq detects drought-responsive *cis*-regulatory loci. **a-b**) Morphological phenotypes of the more tolerant and susceptible B73xOh43 (**a**) and B73xMo18W (**b**) F1s, respectively, grown under well-watered (WW) and drought (DS) conditions. **c**) Allelic distribution of MOA reads at bQTL (B73-chr1:198205029) in the *ZmSO* promoter. The 5 F1s sharing the B73 allele (C/C), were compared to 18 F1s carrying C/A alleles. **d**) Haplotype-specific mRNA counts grouped by the *ZmSO* bQTL in (**c**). Amounts of F1 haplotypes are normalized to the B73 WW level (n=21 F1s which permitted haplotype-specifically RNA analysis). **e**) Allelic distribution of MOA reads at a bQTL (B73-chr2:28118442) in the *ZmTIP3d* promoter. The 12 F1s sharing the B73 allele (C/C), were compared to 13 F1s carrying C/G alleles. **f**) Haplotype-specific mRNA abundance grouped by the *ZmTIP3d* bQTL in (**e**). Haplotype values are normalized to the B73 DS mRNA level (n=23 F1s which permitted haplotype-specifically MOA/RNA analysis). n.d. not detected **g**) Average, normalized MOA coverage for B73xCML333 near *ZmTINY* under WW (blue line) and DS (red line). Roman numeral highlight respective peaks (peak V deletion in B73) and yellow shading indicating significant differences (p<0.05). **h**) Haplotype-specific mRNA counts of *ZmTINY* for B73xCML333 (C333) under WW and DS conditions. **i**) Relative luciferase levels in maize leaf protoplasts harboring either 1.5 kb of the B73 or CML333 proximal promoter allele upstream of the TSS of *ZmTINY* with and without 1-100 µM ABA treatment. **k**) Heatmap of MOA allelic bias at AMP loci under well-watered and drought stress conditions. AMPs (under WW, DS or both) in drought-responsive MOA peaks (p<0.5) are displayed for B73xOh43. Color scale ranging from green (100% bias towards B73) to blue (100% bias to Oh43), gray represents MOA signal below threshold. Clusters represent: (I) allele-specific occupancy in one condition and below detection limit in the other, (II) allele-specific occupancy with consistent bias under both conditions, (III) allele-specific occupancy with bias in opposite direction under the two conditions, and (IV) occupancy with a significant allele-specific bias under only one condition.

To further test the correlation of DS-bQTL and drought-responsive promoter activity we analyzed the maize homolog of the *Arabidopsis* drought-inducible AP2/ERF TF TINY independently in a transient expression assay. The AtTINY family regulates the interplay between plant growth and drought response (Xie *et al*. 2019). Over-expression of *AtTINY* increases drought tolerance at the cost of severely stunted growth, a limitation often observed with drought-related TFs (Xie *et al*. 2019). The maize homolog of *TINY* (Zm00001eb120590) is a candidate gene for drought and leaf size variation (Shikha *et al*. 2017). We found DS-bQTL in multiple MOA footprints surrounding *ZmTINY* which showed significantly higher occupancy in e.g., CML333 and Oh43 compared to B73 under drought (ranging from 1.4 to 5.5-fold higher, Fig. 4c, Additional File 1, Fig. S22). Similarly, higher MOA occupancy under DS for CML333 and Oh43 compared to B73 was also observed downstream of *ZmTINY* (Additional File 1, Fig. S22). These variations in MOA footprints were correlated with allele-specific transcript levels of *ZmTINY*. In F1s under DS, mRNA transcripts of the CML333 and Oh43 alleles were 84-fold and 18-fold more abundant than B73 transcripts, respectively (Fig. 4d, Additional File 1, Fig. S22). MOA signals in the *TINY*^B73^ and *TINY*^CML333^ upstream promoters showed the highest correlation to *ZmTINY* mRNA levels (Additional File 1, Figure S21). We tested these sequences in a dual-luciferase expression assay with and without abscisic acid (ABA) treatment to simulate DS. Both promoter fragments exhibited significantly higher LUC/REN ratio than the vector control. Consistent with trends observed for MOA and mRNA levels (Fig. 4d-e), prom::*TINY*^CML333^ showed higher LUC/REN ratio than prom::*TINY*^B73^ under WW conditions, and exogenous application of 1 and 10 µM ABA further increased the LUC/REN ratio significantly for protoplasts harboring the prom::*TINY*^CML333^ but not prom::*TINY*^B73^ fragment by 41.4% and 60.3%, respectively. Together the results support previous findings of ZmTINY as a drought candidate gene and indicate that bQTL can identify *cis*-elements which act condition-dependent. That said, the drought-responsive regulation of *ZmTINY* may include additional loci, such as the drought-responsive loci downstream.

Differences in *cis*-element coverage between WW and DS conditions at allele-specific sites could be due to changes in the level of occupancy, the direction of allelic-bias, or a combination thereof. To better understand which scenario is more common, we clustered drought-responsive AMPs in the B73xOh43 F1 (31,084 allele-specific MOA variants located in drought responsive footprints). The results showed that more than 80% of drought-responsive AMPs changed MOA occupancy levels between WW and DS, either from no detectable MOA signal to haplotype-specific binding (∼47%, group I), or in the amount MOA coverage between WW and DS conditions while maintaining their allelic bias (∼34%, group II) (Fig. 4f). In contrast, only about 18% of AMPs showed bias changes, either from no significant bias in one condition to a significant bias in the other (∼16%, group IV), or changing the direction of the allelic bias (∼2%, group III) (Fig. 4f). While groups I and IV are somewhat dependent on statistical cut-offs (peak calling and thus AMP definition and/or calling allelic bias), Groups II and III show an allele-specific bias under both conditions. Focusing on Group II and III it becomes evident that changes in allelic bias are ∼15 fold less frequent compared to constant binding bias accompanied by overall changes in MOA-signal. Similar clusters between WW and DS were observed for AMPs in all the 25 F1s (Additional file 1, Fig. S23). We therefore propose that the majority of DS-induced *cis*-element occupancy dynamics at sites of functional genetic variation results from condition-specific TF abundance changes rather than changes of allelic bias between WW and DS conditions.

## Conclusions

We present a robust, high-throughput method for identifying functional, (epi-)genetic variants linked to trait variation in plants. By integrating haplotype-specific TF footprints and transcript abundance, F1 hybrids, and local association mapping at putative *cis*-element loci, we defined a pan-cistrome of the maize leaf under well-watered and drought conditions. Utilization of concatenated dual-reference genomes and F1 hybrid analysis resolved issues of reference bias, *trans*-effects, and technical variation that commonly compromise haplotype-specific quantitation.

Our analysis demonstrates a high level of variation in *cis*-regulatory networks among 25 diverse maize genotypes, and provides a high-resolution map of regulatory elements underpinning the function of nearly two-hundred thousand putative *cis*-element loci in the maize leaf. Finally, we highlight the relevance of bQTL loci for understanding phenotypic diversity in maize, demonstrating that haplotype-specific MOA-seq allowed us to capture the majority of additive genetic variation for most tested phenotypes in maize leaves.

## Author Contributions

T.H. conceived the research project. J.E., S.J.S., J.R.I., D.E.R., M.B.H., B.S., M.St., W.B.F., and T.H. designed experiments. J.E., S.J.S., T.K., V.C.S., A.S., J.V.TR, A.K., and T.H., performed experiments. J.E., S.J.S., A.S., A.K., M-K.B., M.S., J.R.I., D.E.R., G.S., V.C.S, F.F., A.S., A.S.S., J.V.TR, J.C.S., M.B. and T.H. analyzed the data. J.E., S.J.S, A.S.S., and T.H. developed the hybrid mapping pipeline and J.E., S.J.S., J.R.I., D.E.R., J.E., J.V.TR., S.J.S., M.S., A.K., M.B. and T.H. developed analysis scripts. J.E., J.R.I., and T.H. wrote the manuscript, and W.B.F., H.W.B., F.F., S.J.S., and B.S. revised it.

## Acknowledgement

We are grateful to Marty Sachs for germplasm support and the Maize COOP. We thank Martin Krist and Florian Esser for field support. We thank S. Pophaly and M. H. Thoben at MPIPZ for IT support. We thank Christian Becker, Kerstin Becker, and Bruno Huettel for library and sequencing support. We are grateful to Sergius Weizel, Max Blank, Micail D. Mueller, Ram S. Kumar, Guy Koentjes, Isabelle Tol, Sohini Mukherjee, Nafiseh Sargheini, Duong Doan, Luca Watschke, Viola Rajan for greenhouse / experiment support. We are grateful to Stefan Scholten for great help with mite prevention. We would like to thank Felix Andrews for suggesting a mascot for this work, but international restrictions on the transport of fossil bird specimens prohibited its use. We also like to thank Nathan Springer for his critical discussion of the manuscript.

## Data availability statement

All MOA-seq and RNA-seq data discussed in this publication have been deposited at NCBI SRA under accession number PRJNA1101486. Custom scripts have been deposited in the Github repository (https://github.com/Snodgras/MOA_Analysis) under the GNU General Public License v3.0. Other scripts and software used in this study are included in the “Methods” section.

Coverage and binding frequency for all bQTL is accessible via a custom genome browser at https://www.plabipd.de/ceplas/?config=maize_hartwig_config.json&session=local-R7uRvOZKNy9uyYx-ruAIE.

## Materials and Methods

### Plant materials

B73, Mo17, A619, W23, W22, A188, and US-NAM seeds were supplied by the GRIN National Agricultural Library. Seeds were pre-germinated for 48h at 28-30 °C. Each pot contained soil equalized by volume and four seedlings. Plants were grown side by side in greenhouses, under long-day conditions (16h day/8h night, 28-30 °C) for approximately 26 days until 75% of the plants per pot showed the formation of the leaf 4 auricle. Plants were then randomized and 12 pots per treatment were grown either with or without periodic watering through a bottom drench system for 86h. Plants were then harvested and the leaf blades of the oldest leaf without an yet formed auricle were immediately frozen in liquid nitrogen.

### Relative water content and field capacity measurements

At harvest, a 3 cm leaf sheath section below the third leaf ligule was removed for each plant. Four sheath sections (one of each plant in the pot) were processed together to yield one RWC value per pot. First, the fresh weight (FW) of the pieces was determined, followed by determination of the turgor weight (TW) after 24h in water at 4 degrees. The dry weight (DW) was measured after several days at 60 degrees and the relative water content (RWC) was calculated as ((FW-DW)/(TW-DW))*100. Field capacity was measured using a FOM2 Field Operating Meter (ETest, Poland).

### MOA-seq and RNA-seq sample preparation and sequencing

MOA-seq sample and library preparation was performed using 1 g of frozen, finely ground leaf blade tissue following the procedure described in (Savadel *et al*. 2021). RNA-seq of hybrid leaf blade tissue was performed as described in (Hartwig *et al*. 2023).

### MOA-seq data analysis

Reads were filtered using SeqPurge (v2022-07-15) with parameters “-min_len 20 -qcut 0” (Sturm *et al*. 2016). Due to the short fragment length in MOA, read pairs almost completely overlapped. MOA-seq paired-end reads were merged into single-end reads, including base quality score correction, using NGmerge (v0.3) (Gaspar 2018) with parameters “-p 0.2 -m 15 -d -e 30 -z -v”. Diploid genomes were created by concatenating the B73 V5 genome with the respective paternal genome (NAM v1 / 2 genomes (Hufford *et al*. 2021), Mo17 CAU v1, W22 v2 (Springer *et al*. 2018), A188 v1 (Lin *et al*. 2021), and A619 see separate entry). Reads were mapped to the diploid genome (or the separate genomes for inbred data) using STAR (v2.7.7a). As STAR was originally designed to map RNA, we set the flag --alignIntronMax 1 for DNA (no introns allowed) as well as parameters “--outSAMmultNmax 2, --winAnchorMultimapNmax 100, and -outBAMsortingBinsN 5 (Dobin *et al*. 2013).

Bam files were converted to bed format using bamToBed (Quinlan and Hall 2010). Format conversion and the calculation of the average mapped fragment length (AMFL) was done using SAMtools (v1.9) (Li *et al*. 2009). The effective genome size was calculated using unique-kmers.py (https://github.com/dib-lab/khmer/) with AFML and respective genome fasta as inputs.

To generate fragment-center tracks, each mapped read was shortened to 20 bp centered around the middle of the read. Read shortening was performed using awk: for reads with uneven number of bases, the middle base was taken and then read extended 10 bp to each site. For reads with even numbers of bases, one of the two middle bases was chosen randomly and the reads were extended 10 bp to each site. The function genomeCoverageBed of Bedtools (v2.29.0) was then used to convert the bed files to bedgraph, scaled by the quotient of the effective genome size and the number of uniquely mapped reads (reads per genome coverage, RPGC). BigWig files for visualization were generated using bedGraphToBigWig (Kent *et al*. 2010).

### MOA-seq whole genome sequencing control

To avoid potential quantification artifacts due to artificial deviation from the expected 1:1 allele ratio (e.g., read mapping errors), we used equal amounts of whole genome sequencing reads of each haplotype treated and analyzed like MOA reads as control. WGS reads for all lines were downloaded from NCBI SRA (Jiao *et al*. 2017; Springer *et al*. 2018; Lin *et al*. 2021; Hufford *et al*. 2021). All reads were quality trimmed using seqtk (https://github.com/lh3/seqtk) trimfq (v1.3 r106), then cropped to 65 bp length using trimmomatic (v0.39) (Bolger *et al*. 2014) and 600 million reads per haplotype (except B73xIL14H with 300 million reads each) were extracted at random using seqtk sample (seed -s 100). Per F1, B73 and parental reads were merged and mapped as single-end reads to the F1 genome using STAR (2.7.7a) with the same parameters as for MOA reads above. Bam files were converted to bigwig/bedgraph using BamCoverage (v) with the same parameters as for MOA reads above.

### Personalized A619 genome

During this study the maize inbred A619 had no reference assembly. Of the inbred maize lines with high quality assemblies, A619 is most closely related to Oh43 (Flint-Garcia *et al*. 2005). We downloaded approximately 30x coverage of A619 WGS data from NCBI SRA (SRR8997919, SRR10127976, SRR8907067, SRR5725670, SRR5663982, and SRR5663981) and then mapped it to the Oh43 genome using bwa-mem2 (v2.2.1) (Li and Durbin 2009). Only uniquely mapping reads (q30) were retained. The GATK4 pipeline in combination with the “best practices workflow” (https://gatk.broadinstitute.org/hc/en-us/sections/360007226651) was used to identify initial A619/Oh43 SNPs and INDELs with the hard filtering settings: “QD < 2.0”, "QUAL < 100.0", "SOR > 3.0", "FS > 60.0", and "MQ < 40.0". We used these initial variants for base- and variant-recalibration, followed by two additional rounds of training based on convolutional neural networks (CNN) to optimize filtration settings (training input setting: “prior=15.0" and “tranche=99”) and, ultimately selected a refined set of biallelic, homozygous A619/Oh43 SNPs and INDELs. G2Gtools (v. 0.2.7, https://churchill-lab.github.io/g2gtools/) was employed to integrate SNPs and INDELs into the Oh43 genome and generate a A619 pseudo-reference fasta and gff3 file. The average SNP mismatch rate of respective MOA reads from the F1 to the B73xA619 genome was 0.1%, only twice that of the average high-quality B73xNAM genomes (0.04-0.05%), and substantially lower than mapping to either B73 or the A619 psuedo-reference alone (∼0.54%).

### MP and AMP identification

To enable translations between coordinates of the B73 genome and the paternal genomes, hal files were generated using the cactus function of progressive cactus (Armstrong *et al*. 2020) with standard parameters. SNPs between B73 and the paternal lines were determined with the halSnps function of cactus, using parameters “unique” and “noDupes”. From the resulting SNP lists, we selected all SNPs that carried either the B73 or one other base in all analyzed lines (biallelic SNPs,117,898,189). Of the remaining SNPs, only those occurring in at least two of the 25 parental lines were retained (60432443, MAF 0.08). For allele-specific analysis, B73 coordinates of the filtered, biallelic SNPs were translated back to paternal coordinates using halLiftover (hal-release-V2.1) and all SNPs with ambiguous corresponding positions in one of the two parental genomes were removed (de-duplicated biallelic SNPs in at least two lines, 58823746). At each SNP position, we counted RPGC values for both alleles using bedtools map (bedtools version v2.29.0) and calculated the read numbers corresponding to the RPGC numbers for further calculation. Binding frequencies (BF) at SNP positions were determined as RPGC-B73/(RPGC-B73 + RPGC-Pat). For MPs we retained WGA SNPs that were located within MOA-peak, and had more than 7 RPGC (>∼25 reads) for at least one allele combined with at least one read in the corresponding allele. Allele specific binding at MPs (significant deviation from the expected 0.5 BF in the case of equal binding) was determined via binomial testing in R 4.1.1. SNP positions with a false discovery rate corrected p-value <0.01 were considered allele-specific MPs (AMPs). Additionally, we determined the allelic ratio of WGS control reads in a 65bp window at all MP and excluded all AMPs with a WGS ratio above or below the upper and lower 5 percentile value of all MPs, respectively.

### Peak calling

For peak calling MOA bam files were used with MACS3 (v3.0.0a7, https://github.com/macs3-project/MACS) using the following parameter: -s and --min-length “AMFL”, --max-gap “2x AMFL”, –nomodel, –extsize “AMFL”, --keep-dup all, -g “effective genome size”. AMFL represents “Average MOA Fragment Length” calculated with samtools stats at default parameters.

### Treatment-specific peak calling

MOA-alignment bam files were converted to bed format using bedtools bamToBed (v2.29.0). The genomeCoverage function of bedtools was used to convert pooled replicated bed files to bedgraph with the reads per million (RPM) scaling factor. The RPM-normalized coverage difference between treatments was calculated with use of intersect and subtract functions of bedtools. The resulting differences in coverage counts for WW/DS treatment were used to create an unbinned (1 bp resolution) bed file to produce a bigwig coverage track, which was used as input for MACS3 (v3.0.0a7) peak calling using parameters: --min-length 30, --max-gap 60, – nomodel, –extsize 1, --keep-dup all, -g “effective genome size”, -q 0.01. Significant differences between WW and DS peaks were determined by Welch t-test using the individual bio replicate coverages and peaks with a p-value < 0.05 were retained.

### DNA methylation analysis

Parental DNA methylation data of the NAM lines (Hufford *et al*. 2021) was obtained from iPlant (/iplant/home/maizegdb/maizegdb/NAM_PROJECT_JBROWSE_AND_ANALYSES).

Methylation data for non-NAM lines (Liang *et al*. 2021; Lin *et al*. 2021) were obtained as SRA archives (Bioprojects PRJNA657677 and PRJNA635654) and processed as described in Hartwig *et al*. (2023). B73 x Mo17 hybrid methylation data was previously published in Hartwig *et al*. (2023) and showed strong correlation with parental methylation at B73 x Mo17 AMPs (Additional file 1, Fig. S7). Context specific methylation around AMPs/MPs was determined as described in Hartwig et al. (2023), significant differences in DNA methylation were determined following Regulski *et al*. (2013) (one allele <10% methylated and the other >70%). Sites for testing consistency of DNA methylation/ haplotype specific binding relations were selected based on having at least two F1 lines differentially methylated, at least two F1 lines equally methylated, and at least one F1 line AMP at the given site. In this analysis (Fig. 3c), a more stringent definition of equal methylation (as opposed to not being differentially methylated) was employed: Equal methylation was defined as both alleles <10% or both >70% methylated.

### Local association mapping to map bQTL

The binding ratio of the MOA peaks, CH, CHG and CHH binding, and the read depth were collected separately for all hybrid lines for the well-watered and drought-stressed conditions. The binding frequency at loci with no reads was set to NA. Genotyping information and the methylation ratio information for _m_CG, _m_CHG, and _m_CHH were used together with the read depth to conduct local association studies. All MPs with the respective haplotype-specific MOA coverage (binding frequency) and average surrounding (+/- 20 bp) methylation ratios were sampled and used to run five different mixed linear models for each MP, where:

MOApeak=Marker_i_ + CG + CHG + CHH + Readdepth

MOApeak=Marker_i_ + CHH + Readdepth

MOApeak=Marker_i_ + CG + Readdepth

MOApeak=Marker_i_ + CHG + Readdepth

MOApeak=Marker_i_ + Readdepth

All factors, except the read depth, were used as fixed effects. Associated MPs at a false discovery rate of 5% were selected to calculate the explained variance for all five models separately. The explained variance and probability values for each MP were compared across the five models genome-wide to identify any superior prediction ability associated with incorporating any methylation information. The analyses were performed in Julia 1.8.1 and R 4.1.2. The built-in linear model in R and the mixed linear model from the R package lme4 were utilized to estimate predictions of the MOA peaks. bQTL located within 65 bases were clumped into linkage groups (lowest p-value determined lead bQTL).

### RNA-seq analysis

RNA-seq data was mapped to the concatenated diploid genomes using STAR (v2.7.7a), with options --outSAMmultNmax 1, --outFilterMultimapNmax 1, --winAnchorMultimapNmax 100, -- twopassMode Basic, --outFilterIntronMotifs RemoveNoncanonical, --outFilterType BySJout, -- quantMode GeneCounts and using concatenated gff3 file containing gene models from both parents. To determine allele-specific transcript abundance, for each line, B73 and corresponding paternal positions for all SNPs determined by halSnps were generated via halLiftover. Of the resulting position pairs, ambiguous ones were removed. Each SNP was then assigned the B73 genes it overlaps with. The respective NAM gene info was added using a Pan-gene file (downloaded from MaizGDB), retaining strand information in both cases. Mapped read information was converted into read bed files using bamToBed and each SNP was assigned all reads overlapping with it in B73 and at the parental genome coordinates (strand-specific, separately for the three replicates). Only SNPs carrying reads in both alleles were retained to ensure that the SNP was truly located within the gene in both alleles. Afterwards, reads for each gene were counted per replicate (reads which had two or more SNPs were counted only once) and allele. For A188, where no Pan-gene entries were available, SNPs were also mapped onto the A188 gff3 and gene pairs were generated based on shared SNP positions. This way, B73 reads and paternal reads could be compared for differential transcript abundance analysis in DEseq2 (Love *et al*. 2014). Genes with a false discovery rate corrected p-value < 0.05 were considered allele-specific in their transcript abundance.

### Variance Component Analysis Pipeline (VCAP)

To run the Variance Component Analysis Pipeline (VCAP), we required three datasets: (1) genome-wide markers across the Nested Association Mapping (NAM) population Recombinant Inbred Lines (RILs), (2) trait values across NAM RILs for each trait analyzed, and (3) coordinates for MOA peaks or bQTL SNPs across founder lines to partition each component. For the genome-wide markers, we used publicly available resequencing SNPs from the NAM founders (Hufford *et al*. 2021) that had been projected onto the NAM RILs (Cyverse:/iplant/home/shared/NAM/Misc/NAM-SV-projected-V8/). Trait data collected from the NAM RILs (n=143) were curated from publications (Supplemental file NAM_phenotype_metadata.xlsx). We used two sampling schemes to create our MOA partitions. First, we randomly sampled 1 SNP from each MOA peak called from plants under well-watered conditions, resulting in ∼1 million SNPs. Second, only the bQTL SNPs with significant association to the genotype, not methylation, were used to represent MOA. Given many of the significant bQTL were in close proximity and thus linkage disequilibrium with each other, bQTL SNPs were clumped using plink (Purcell *et al*. 2007) within 150 bp of each other and false discovery rate cutoffs at 10% and 30%, resulting in 145,425 SNPs. Any SNP outside of a MOA peak created the non-MOA pool from which the background SNPs were drawn. Because of the non-random distribution of MOA peaks throughout the genome, we also included a matched background component: each MOA SNP was matched to a random non-MOA SNP by allele frequency (number of lines containing the alt allele/total lines without missing data at that position, 0.1 bin size) and distance from the nearest gene (TSS or TTS).

This sampling and matching of MOA and background SNPs created the set of SNPs for estimating a set of kinship matrices for a single VCAP run. Kinship matrices were created for the MOA SNPs, BKGD SNPs, and the rest of the genome (remaining non-MOA and non-BKGD SNPs) using Tassel 5 (Bradbury *et al*. 2007). Under each sampling method (1 SNP per peak or bQTL strict), we sampled 100 times, creating 100 permutations of kinship matrix sets. To calculate heritabilities of all 143 traits, each set of kinship matrices and traits were run through a REML model using LDAK (Speed *et al*. 2012). Thus, the permutations gave us a range of heritability estimates that could result from these particular components, traits, and the population (Figure 3). For the strictly bQTL SNP runs, the same bQTL SNPs were used in every permutation while the background SNPs differed across permutations.

To evaluate the reliability of our heritability estimation method, we simulated traits with defined contributions from specific sets of kinship matrices and compared estimates of the heritabilities generated by the above VCAP protocol. We used one of our previously generated kinship matrix sets (1 SNP per peak sampling) to simulate traits assigned certain heritabilities for each component (four sets of heritabilities, 10 traits per set). We simulated traits as the sum of four normally distributed random vectors, each with zero mean and covariance equal to one of the three kinship matrices or the identity matrix (for residual variation) multiplied by a specific heritability value. The simulated traits and kinship matrices were used in the REML modeling step to estimate heritabilities. Estimated heritabilities were then compared to known heritabilities. All scripts written for the analyses in the study were deposited at https://github.com/Snodgras/MOA_Analysis.

### MOA bQTL and eQTL linkage analysis

Linkage disequilibrium was calculated between the binding QTL reported in this study and a set of 10,618 cis-eQTL identified based on expression data the roots of 340 maize genotypes (Sun *et al*. 2023). Genomic coordinates of the 98,383 binding QTL on the B73_RefGen_V5 maize genome were converted to B73_RefGen_V4 positions using CrossMap as implemented in EnsemblPlants (Zhao *et al*. 2014; Howe *et al*. 2020). 99.5% of binding QTL positions could be successfully converted to B73_RefGen_V4 positions and, of these, 36,777 were present in a set of 12,191,984 genetic markers segregating in the population of 340 maize lines used to conduct eQTL analysis with a minor allele frequency ≥ 0.05 and less than 2% of genotypes exhibiting heterozygous genotype calls. Linkage disequilibrium was calculated between bQTL markers and cis-eQTL markers in all cases where a cis-eQTL and a bQTL were located within ten kilobases of each other using genotype calls from the 340 maize varieties (Purcell *et al*. 2007; Sun *et al*. 2023). To assemble a control set of genetic markers with the same properties as the bQTL, bQTL that were successfully converted to B73_RefGen_v4 and matched to Sun et *al*. markers were divided into ten bins based on their distance from the closest annotated transcription start site (0-1 kilobase, 1-2 kilobases, and so on), plus two additional categories for intragenic SNPs and SNPs > 10 kilobases from the nearest annotated gene. A random subset of 2 million B73_RefGen_v5 SNPs used to detect bQTL were also converted to B73_RefGen_v4 and matched to segregating markers from Sun et *al*., as described above. These markers were subsampled to create a second set of 36,777 control markers with representation in each of the twelve bins equal to the levels observed for the real bQTL.

### Further data processing

To obtain the high confidence list of drought-responsive MOA regions, all MOA bQTL (unclumped WW or DS associated SNPs (FDR=0.5), were filtered for overlap with AMPs located in regions with significantly (p<0.05) increased or reduced MOA occupancy between DS and WW conditions in at least two F1s. The retained 3198 and 13335 drought induced and repressed, respectively, MOA loci,.

Comparisons and calculations of lists were either performed in bedtools intersect or custom awk and bash scripts. Hypergeometric tests for over/under-representation and data visualization were performed in R. Pearson correlation coefficients of bigwig file format MOA-seq data were calculated and visualized using the multiBigwigSummary and plotCorrelation functions of deepTools (Ramírez *et al*. 2016) with a window size of 1000 bases.

