## Additional File1, Supplemental Figures1-23 for "Genetic variation at transcription factor binding sites largely explains phenotypic heritability in maize"

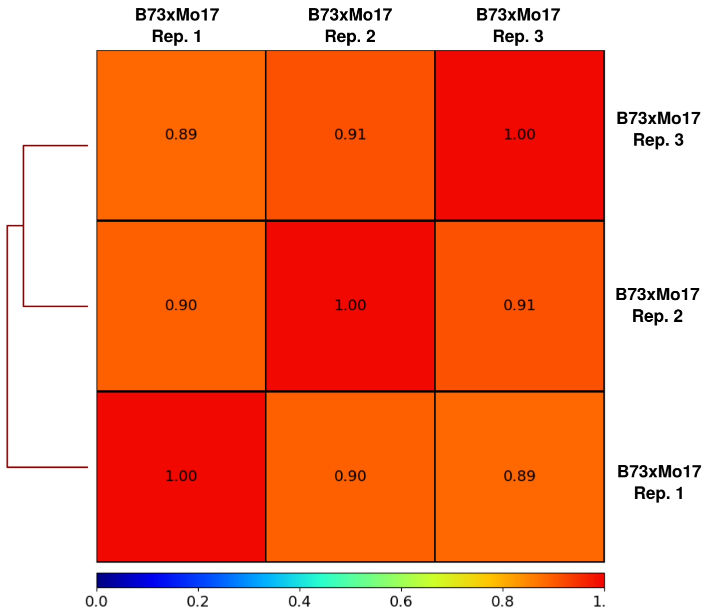


Figure S 1: Pearson correlation coefficients for MOA-seq replicates of B73xMo17.

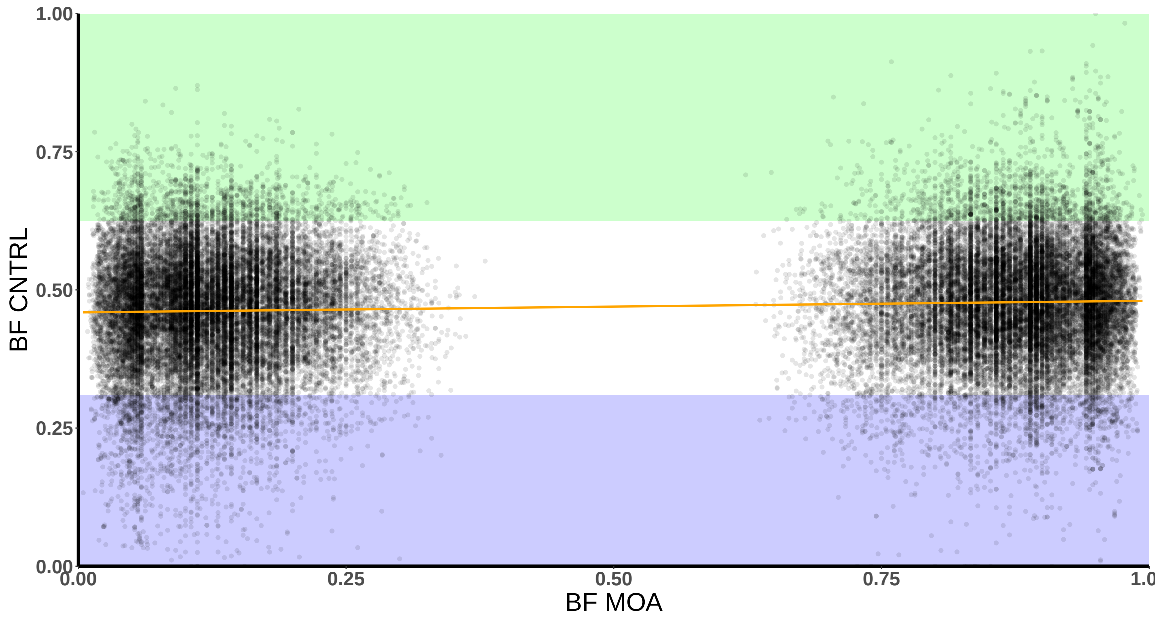


Figure S 2: Visualization of AMP correction via whole genome sequencing control.

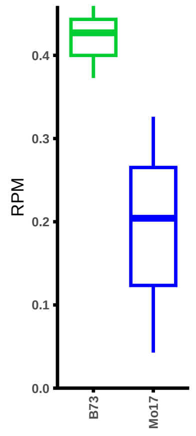


Figure S 3: Allele-specific mRNA abundance of *ZmBIF2*.

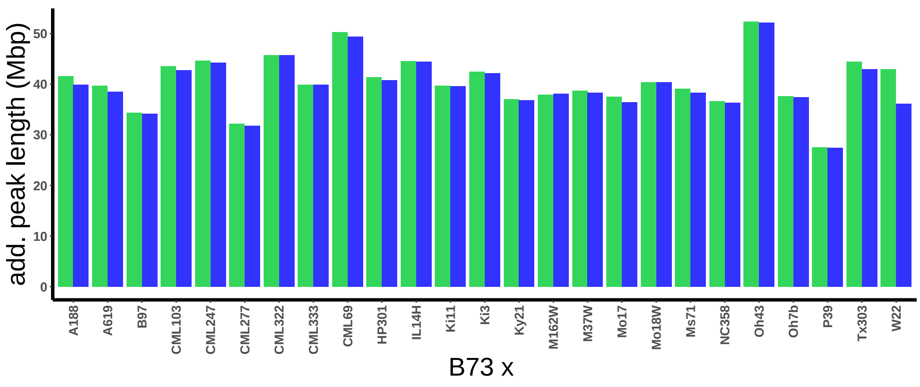


Figure S 4: Bases covered by MOA-peaks in each hybrid for the B73 genome fraction (green) and the NAM genome fraction (blue) in Megabases.

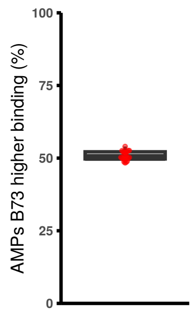


Figure S 5: Fraction of AMPs with higher binding towards the B73 allele in the 25 hybrids.

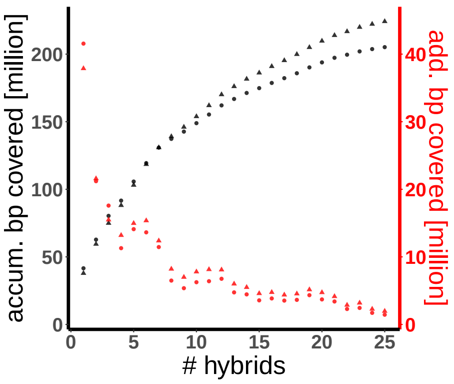


Figure S 6: Cumulative number of bp covered by MOA-seq peaks in the 25 F1 hybrids (black) and number of base pairs added by including each hybrid (red).

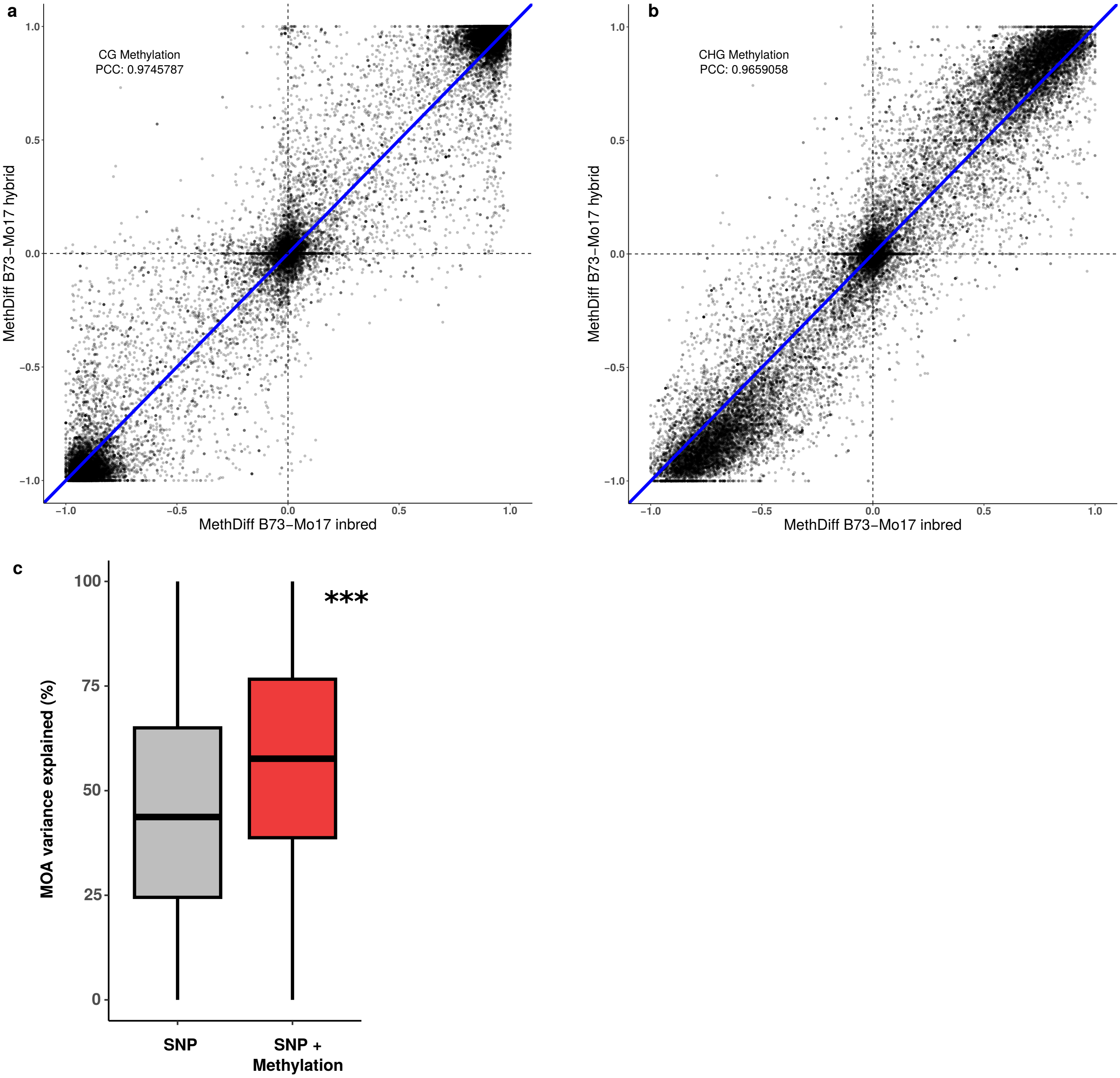


Figure S 7: Correlation of DNA methylation at AMP loci between F1 and parental alleles and TF footprint variation explained by genotype and methylation.

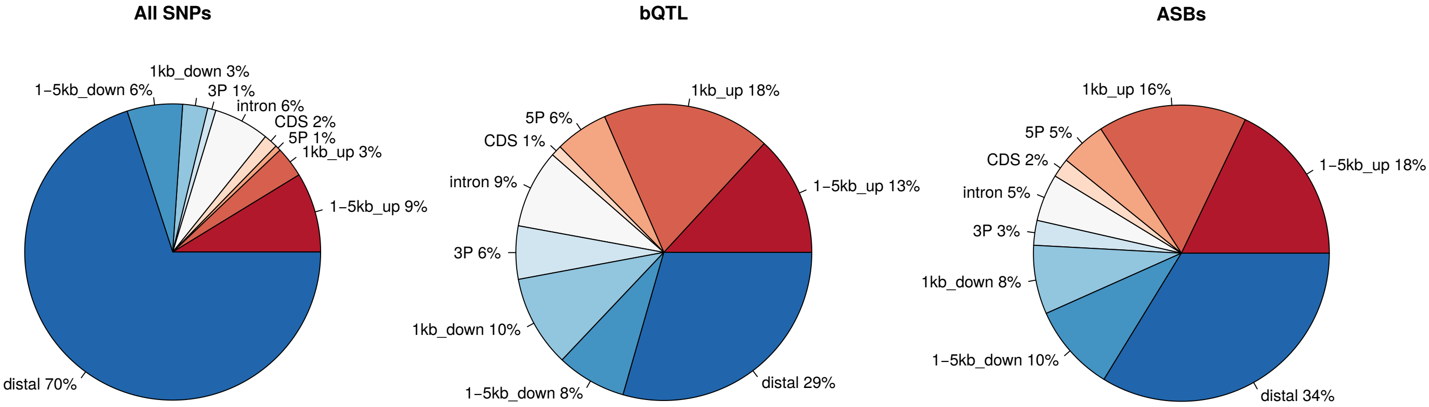


Figure S 8: Pie charts displaying the distribution of SNP positions relative to genes.

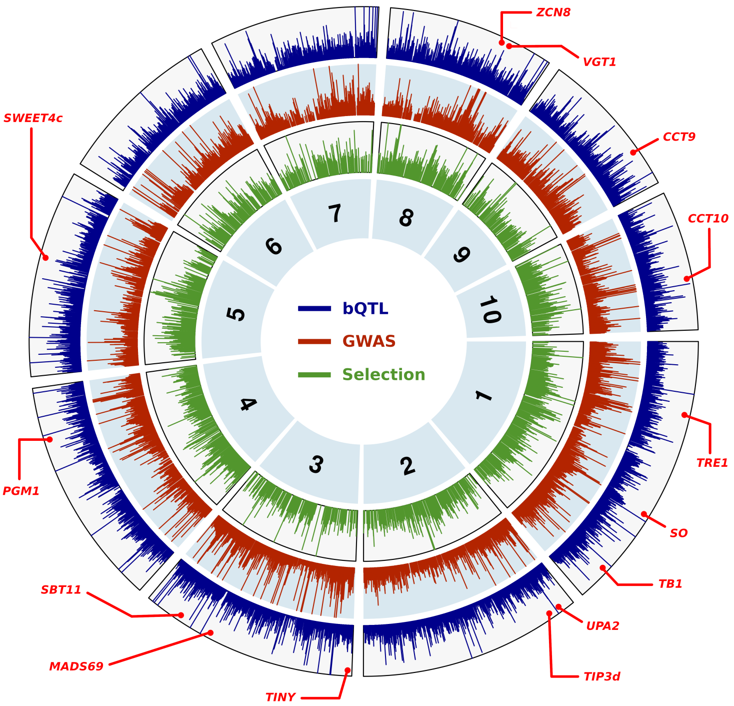


Figure S 9: Distributions of bQTL, GWAS hits, and selective sweep genomic features across ten maize chromosomes.

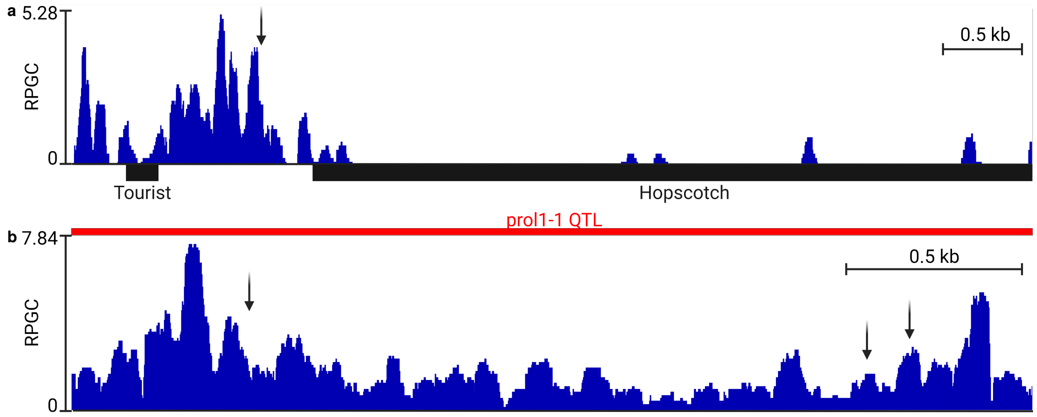


Figure S 10: Genomebrowser view of bQTL regions of *ZmTB1* (a) and *ZmGT1* (b).

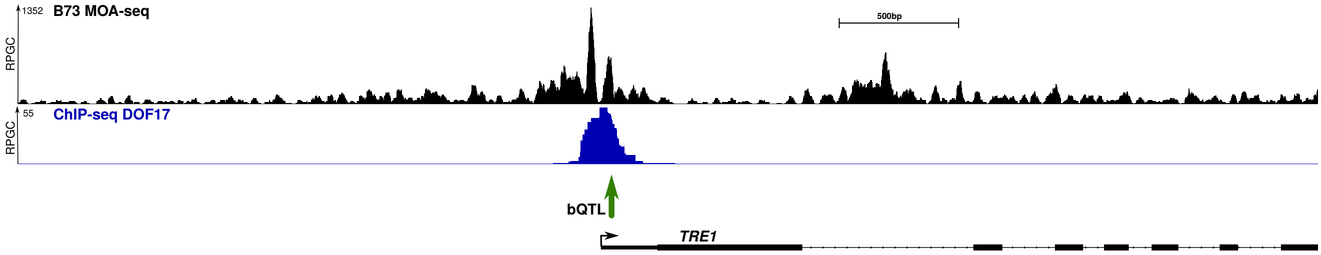


Figure S 11: Genomebrowser view of bQTL (green arrow) in the *ZmTRE1* promoter with B73 MOA high-resolution track and ZmDOF17 binding site (Tu et al. 2020).

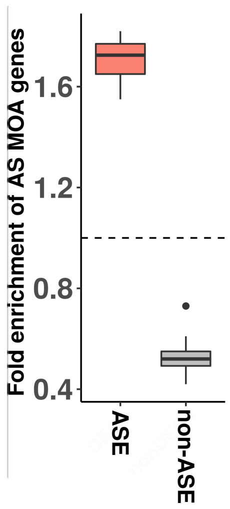


Figure S 12: ASE and non-ASE genes under DS conditions are significantly more and less enriched for AMPs in their 3 kb promoter upstream of the TSS, respectively.

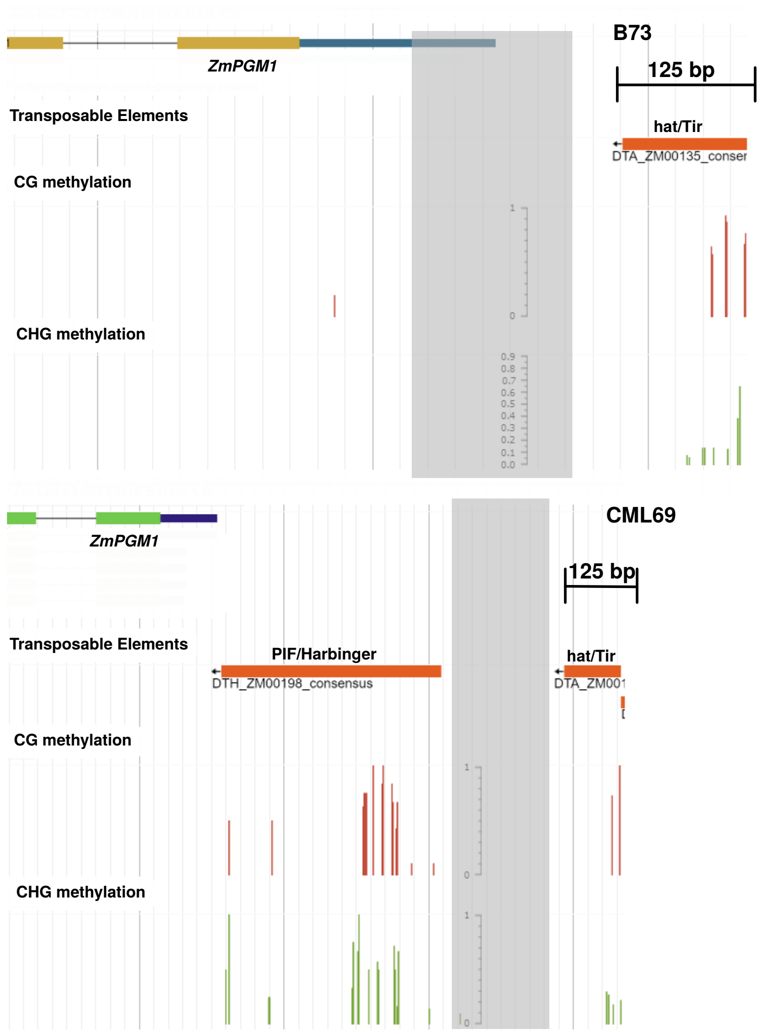


Figure S 13: Genome browser view *ZmPGM1* promoter alleles in B73 and CML69.

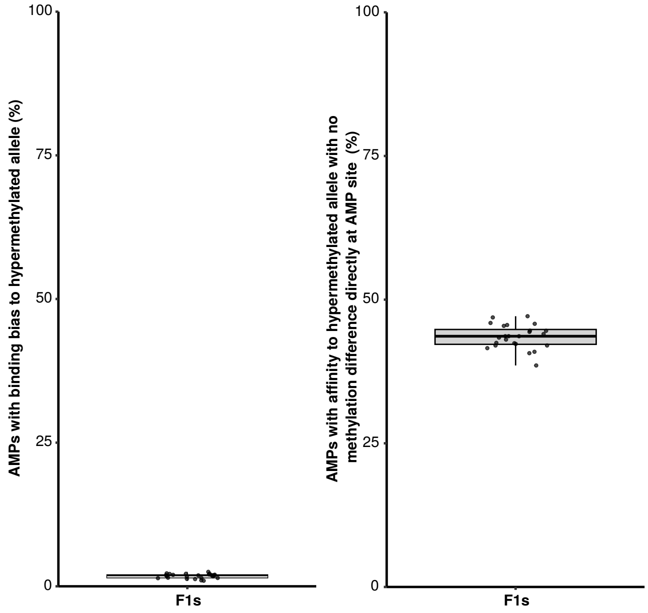


Figure S 14: Additional analysis of AMPs with binding bias towards hypermethylated alleles.

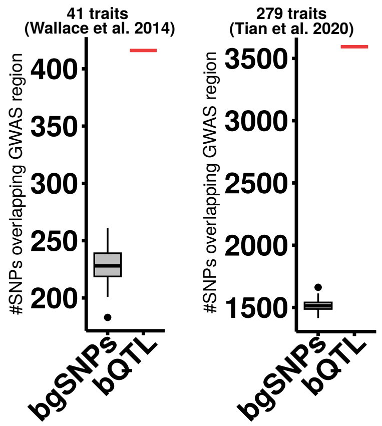


Figure S 15: bQTL are enriched in GWAS hits.

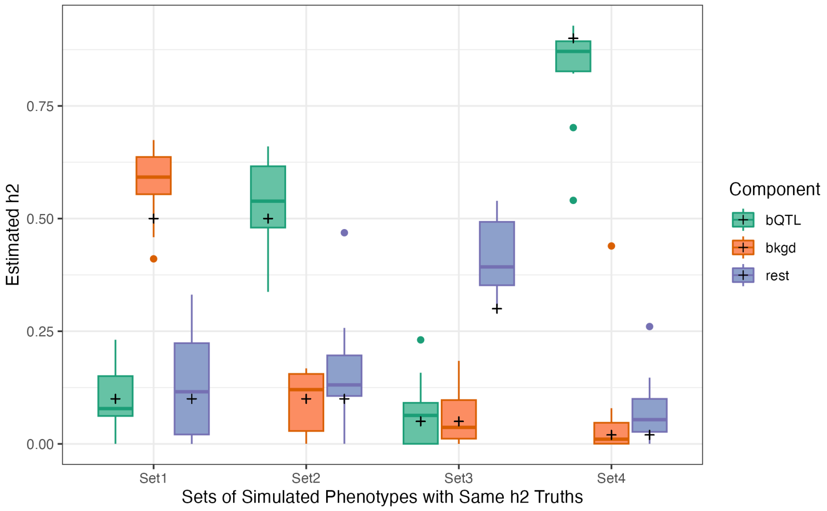


Figure S 16: Simulated trait heritabilities.

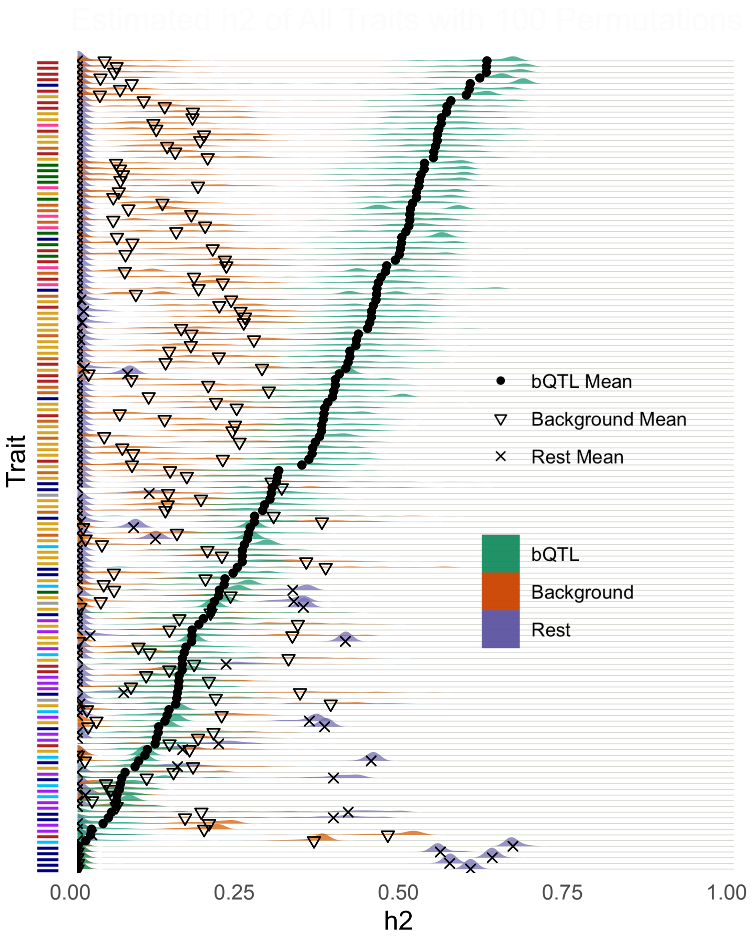


Figure S 17: Estimated additive heritability organized by 143 traits.

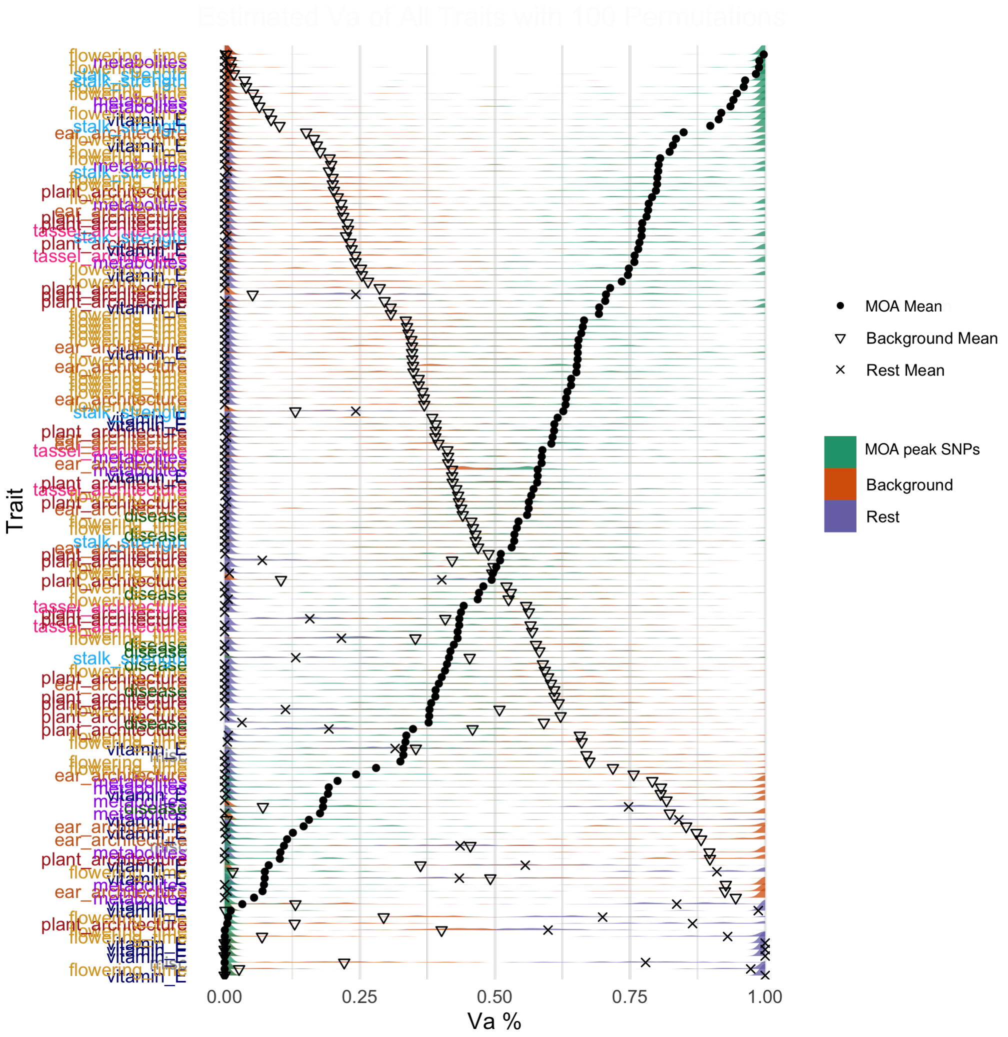


Figure S 18: Estimated additive genetic variance for random SNPs in MOA peaks organized by 143 traits.

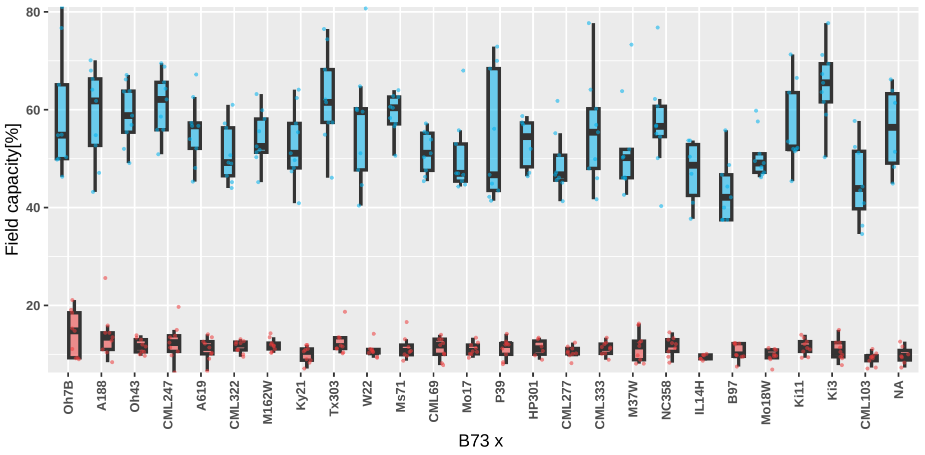


Figure S 19: Field capacity in pots at the moment of harvest.

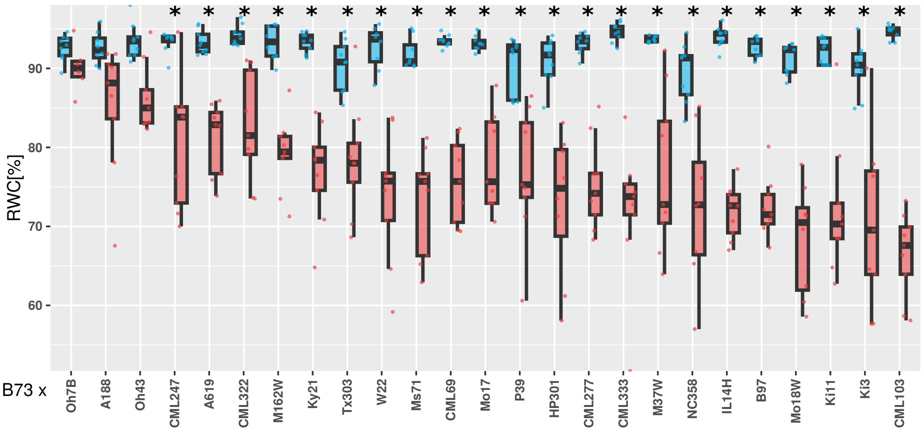


Figure S 20: Assessment of relative water content of the 25 F1 hybrids under well-watered and drought conditions.

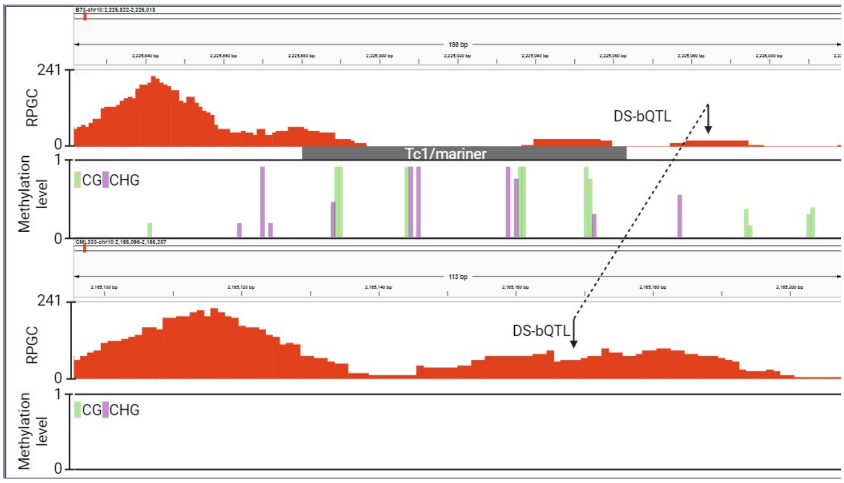


Figure S 21: Genome Browser view of MOA-seq and DNA methylation levels near the causative TE insertion in the *ZmNAC111* promoter under drought conditions.

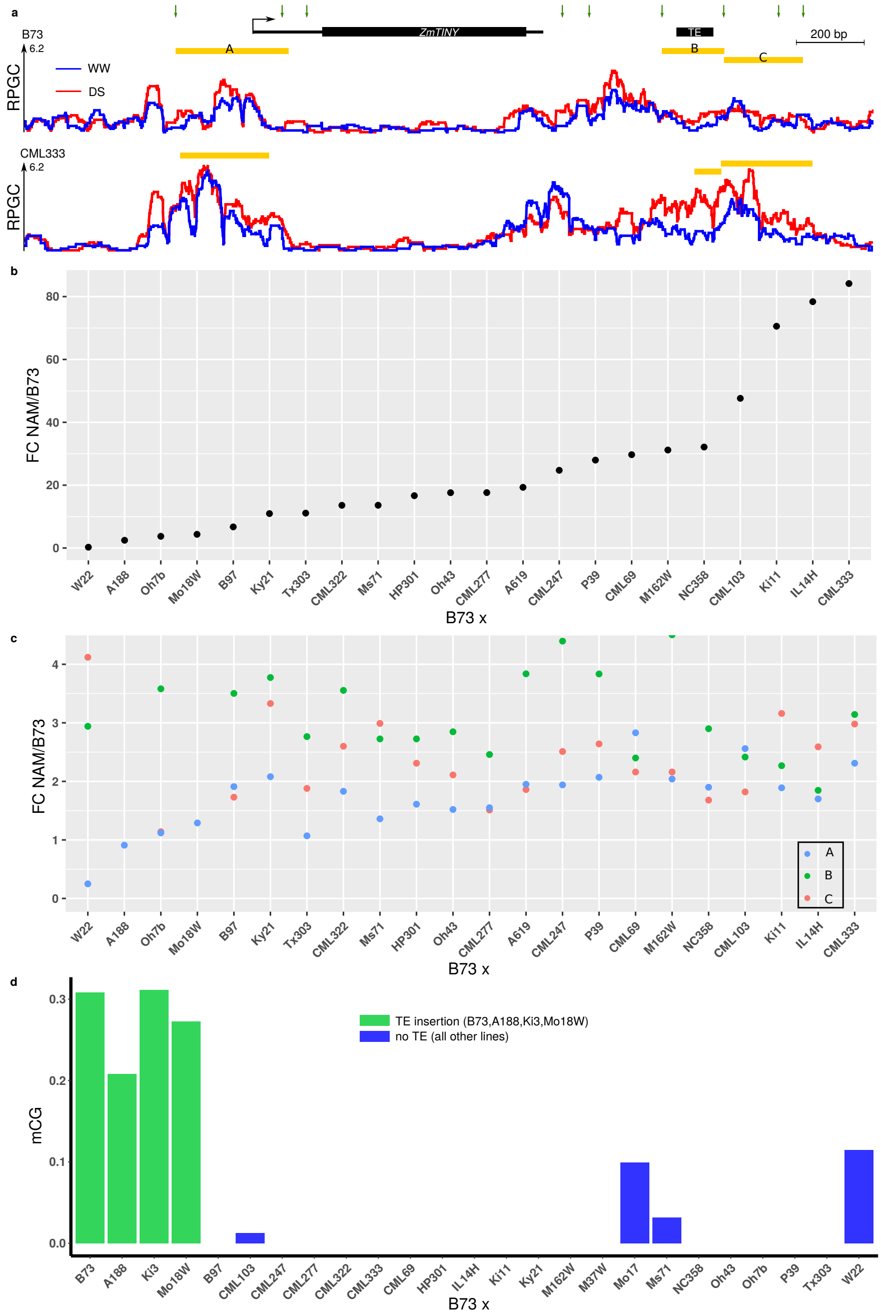


Figure S 22: Overview of the *ZmTINY* locus with allele-specific mRNA abundance and TF-binding.

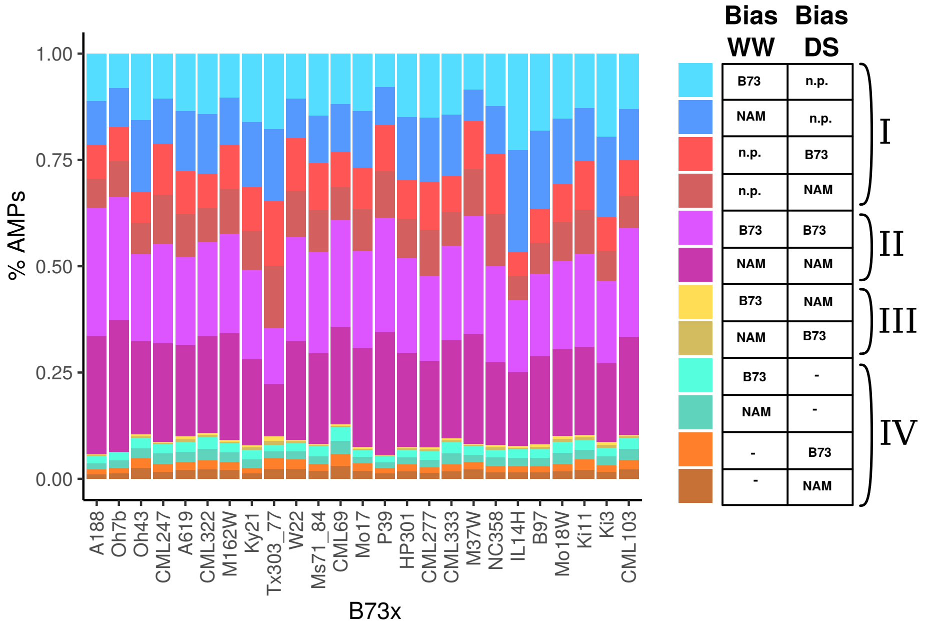


Figure S 23: Comparison of haplotype-specific binding between WW and DS.
