## Supplemental Figure and Table Legends for "Genetic variation at transcription factor binding sites largely explains phenotypic heritability in maize"

**Additional file 1, Supplemental Figure legends**

Figure S1 (Pearson correlation Mo17 replicates MOA): **Pearson correlation coefficients for MOA-seq replicates of B73xMo17.**

Figure S2 (WGS control): **Visualization of AMP correction via whole genome sequencing control.** Each dot represents one AMP, shaded areas indicate the fraction of AMPs excluded due to bias in counts towards one allele in the control data. The orange line represents a linear regression.

Figure S3: **Allele-specific mRNA abundance of *ZmBIF2* transcripts.** RPM: reads per million.

Figure S4: **Bases covered by MOA-peaks in each hybrid for the B73 genome fraction (green) and the NAM genome fraction (blue) in Megabases**.

Figure S5: **Fraction of AMPs with higher binding towards the B73 allele in the 25 hybrids.**

Figure S6: **Cumulative number of bp covered by MOA-seq peaks in the 25 F1 hybrids (black) and number of base pairs added by each consecutive hybrid (red).** B73 parts (circles) for each hybrid and NAM parts (triangles) for each hybrid genome are depicted separately. Peak coordinates for the NAM parts of each genome were projected to the B73 genome to allow this comparison.

Figure S7: **Correlation of DNA methylation at AMP loci between F1 and parental alleles and TF footprint variation explained by genotype and methylation. a** and **b:** Methylation differences are expressed as Mo17 methylation values subtracted from B73 methylation values in inbred and hybrid comparisons (a: CG, b: CHG). The blue line indicates x=y for orientation. PCC: Pearson Correlation Coefficient.
**c**: Percentage of MOA-seq variation explained by genotype alone (red) and genotype combined with DNA methylation (gray). *** indicates a significant difference (Wilcoxon Signed-Rank Test, p-value < 2.2e-16)

Figure S8: **Pie charts displaying the distribution of SNP positions relative to genes.** All SNPs are all SNPs tested in this study, i.e. all biallelic, one-to-one mappable SNPs occurring in at least two lines and bQTL are WW bQTL, ABS are allele-specific binding sites of ZmBZR1 (Hartwig et al. 2023).

Figure S9: **Distributions of bQTL, GWAS hits, and selective sweep genomic features across ten maize chromosomes.** From outer to inner circles tracks are as follows: ① chromosome names, ② selective sweeps (Xu et al. 2020) detected between modern maize and teosinte, ③ p-values of the maize GWAS Atlas lead SNPs of leaf/whole plant traits (Lian et al. 2020), and ④ bQTL associated with variation in MOA coverage in the pan-cistrome. Red markers denote bQTL near classical domestication and flowering time genes.

Figure S10: **Genomebrowser view of bQTL regions of *ZmTB1* (a) and *ZmGT1* (b).** bQTL locations are indicated with vertical black arrows.

Figure S11: **Genomebrowser view of bQTL (green arrow) in the *ZmTRE1* promoter with B73 MOA high-resolution track and ZmDOF17 binding site (Tu *et al.* 2020).**

Figure S12: **ASE and non-ASE genes under DS conditions are significantly more and less enriched for AMPs in their 3 kb promoter upstream of the TSS, respectively.**

Figure S13 **Genome browser view *ZmPGM1* promoter alleles in B73 and CML69.** CG and CHG DNA methylation levels and the TE insertion sites (PIF TE only in CML69) for both haplotypes are shown. The position of the MOA peak from Fig. 2i is highlighted.

Figure S14 **Additional analysis of AMPs with binding bias towards hypermethylated alleles.** Left panel: Percentages of AMPs with a stronger binding to the hypermethylated allele for each hybrid. Right panel: Percentages of hypermethylated (in +-20bp around the AMP), bound AMPs that display no differential methylation in +- 5bp surrounding the AMP.

Figure S15 **bQTL are enriched in GWAS hits.** Enrichment GWAS hits (lead SNP +/- 100 bp) among bQTL compared to 100 bootstrapped sets of matched background SNPs across two curated datasets of 41 and 279 traits (Wallace *et al.* 2014; Tian *et al.* 2020).

Figure S16: **Simulated trait heritabilities.** 10 traits were simulated per 4 sets of heritability scenarios of each component (bQTL, background, rest) and then ran through the VCAP. Estimates for the 10 traits' heritability are shown as boxplots, color by component. The black cross represents the known heritability of the traits simulated for a given set.

Figure S17: **Estimated additive heritability organized by 143 traits.** Colored ridges show the estimated additive genetic variance across 100 permutations for either MOA bQTL (green), bgSNPs (orange), and rest (purple) components. Black symbols represent the mean estimated value across permutations. Traits arranged by bQTL mean-variance estimates and color-coded according to general trait groupings: vitamin E metabolites = navy blue, metabolites = purple, stalk strength = light blue, flowering time = gold, plant architecture = red, disease = green, tassel architecture = pink, ear architecture = orange, misc. = grey.

Figure S18: **Estimated additive genetic variance for random SNPs in MOA peaks organized by 143 traits**. Colored ridges show the estimated additive genetic variance across 100 permutations for either MOA peaks (green, one SNP sampled per peak), bgSNPs (orange), and rest (purple) components. Black symbols represent the mean estimated value across permutations. Traits arranged by bQTL mean-variance estimates and color-coded according to general trait groupings: vitamin E metabolites = navy blue, metabolites = purple, stalk strength = light blue, flowering time = gold, plant architecture = red, disease = green, tassel architecture = pink, ear architecture = orange, misc. = grey.

Figure S19: **Field capacity in pots at the moment of harvest.** Field capacity was measured at harvest for both WW and DS plants.

Figure S20: **Assessment of relative water content of the 25 F1 hybrids under well-watered and drought conditions.** Relative water content at the time of harvest for WW and DS plants. * indicate a significant difference between WW and DS based on ANOVA followed by Tukey post hoc test, $p<0.01$.

Figure S21: **Genome Browser view of MOA-seq and DNA methylation levels near the causative TE insertion in the *ZmNAC111* promoter under drought conditions.** This region carries a Tc1/mariner TE in B73 (upper panel) but not CML333 (lower panel). The bQTL is indicated by a vertical black arrow in both genomes.

Figure S22: **Overview of the *ZmTINY* locus with allele-specific mRNA abundance and TF-binding. a**: Genome browser view of the upstream and downstream region of *ZmTINY* in the hybrid of B73xCML333. Green arrows mark DS bQTL positions. Yellow blocks mark regions analyzed in c. The black TE block marks a TE present in B73, A188, Ki3, and Mo18W. **b**: Fold change (FC) in mRNA abundance between the NAM and the B73 allele under DS conditions. **c**: FC in MOA occupancy between the NAM and the B73 allele under DS conditions. Average MOA occupancy per base was calculated in the three yellow regions marked in a. To ensure a comparison of homologous regions, regions between two homologous SNPs determined by whole genome alignment were chosen. Note that the W22 promoter contains a TE in the A region, adding almost 5000 bases to the W22 region. To ensure this is not the reason for the reduced binding, two more regions in W22 were compared to the B73 A region: only the region up to the TE (FC 0.36) and a region of similar length to the B73 A region (FC 0.21). The downstream regions of A188 and Mo18W do not differ from B73 thus no FC could be detected. **d**: CG methylation in 40bp surrounding SNP B73-chr3:4729798 (directly adjacent to TE downstream of ZmTINY).

Figure 23: **Comparison of haplotype-specific binding between WW and DS.** All AMPs are categorized according to their binding bias in WW and DS. Binding frequencies >=0.6 and <=0.4 were considered biased. The Roman numerals indicate the same four categories displayed in Fig.4 k for the Oh43 hybrid - I: Bias in one condition but no binding in the other; 2:Bias in the same direction in both conditions; 3: Bias in opposite directions; 4:Bias in one condition but no bias in the other condition.

**Additional files 2-13, Supplemental Tables legends:**

Additional file 2, Supplemental Table S1: **Numbers of reads uniquely mapped and peaks detected in the hybrid genomes.**

Additional file 3, Supplemental Table S2: **Genes with MOA peaks within 5kb upstream to 1kb downstream of genes annotated in the concatenated hybrid genome**.

Additional file 4, Supplemental Table S3: **26 parental lines were used to generate hybrids in this study.**

Additional file 5, Supplemental Table S4: **Numbers of reads mapped uniquely to the hybrid genome in each F1 hybrid, per replicate.**

Additional file 6, Supplemental Table S5: **SNPs with significant allele-specific binding bias in 25 hybrids**. All MPs denote the number of SNP in MOA-seq peaks where at least one allele had an equivalent of 25 reads (for details see methods). AMPs are all MPs with a significant deviation from the expected 1:1 ratio determined by binomial testing (FDR corrected p-value < 0.01) with those removed that showed a significant bias (p<0.05) in whole genome sequencing control. “B73 and NAM high affinity” shows the subfractions of AMPs biased to each parent.

Additional file 7, Supplemental Table S6: **Genome coordinates of bQTL under WW conditions with Variance explained and FDR corrected p-values for either the genotype or the three methylation contexts.**

Additional file 8, Supplemental Table S7: **Overlap of ASEs and AMP genes.** Sheet 1: AMP genes are defined as genes carrying an AMP within 3kb upstream of their TSS. ASEs are genes displaying allele-specific mRNA abundance as determined by DEseq2 (p<0.05), non-ASEs are defined as genes with a p-value above 0.95. Enrichment and under-enrichment p-values provided are based on a hypergeometric test. Sheets 2 and 3: ASE genes in WW and DS conditions.

Additional file 9, Supplemental Table S8: **Heritability estimates descriptive statistics across VCAP components**. Descriptive statistics of all heritability estimates of the 100 permutations of VCAP for each trait. Mean, median, and standard deviation of estimated heritability for the bQTL, background, and rest components as well as the total heritability are included.

Additional file 10, Supplemental Table S9: **Number of peaks with significant differential occupancy between WW and DS conditions in the F1 hybrids.**

Additional file 11, Supplemental Table S10: **Genome coordinates of bQTL under DS conditions with Variance explained and FDR corrected p-values for either the genotype or the three methylation contexts.**

Additional file 12, Supplemental Table S11: **High confidence drought candidate genes.** Candidate genes were selected to have allele-specific expression, differential expression between WW and DS, and an overlap with a drought-responsive MOA peak (either more or less binding under MOA) and allele-specific binding in at least two lines (features within a region from 5kb upstream to 1kb downstream of the gene body were considered). Further selection was performed by overlapping with QTLs/eQTLs related to drought treatment (Li et al. 2016; Shikha et al. 2017; Wu et al. 2021).

Additional file 13, Supplemental Table S12: **Phenotype Metadata used in VCAP**. Included are the publication details for every trait used in the VCAP as well as descriptors of the tissue and mapping population used to measure phenotypes and type of phenotype data reported (i.e., blup, blue).
